# Identification of novel inner membrane complex and apical annuli proteins of the malaria parasite *Plasmodium falciparum*

**DOI:** 10.1101/2021.02.03.428885

**Authors:** Jan Stephan Wichers, Juliane Wunderlich, Dorothee Heincke, Samuel Pazicky, Jan Strauss, Marius Schmitt, Jessica Kimmel, Louisa Wilcke, Sarah Scharf, Heidrun von Thien, Paul-Christian Burda, Tobias Spielmann, Christian Löw, Michael Filarsky, Anna Bachmann, Tim W. Gilberger

## Abstract

The inner membrane complex (IMC) is a defining feature of apicomplexan parasites, which confers stability and shape to the cell, functions as a scaffolding compartment during the formation of daughter cells and plays an important role in motility and invasion during different life cycle stages of these single celled organisms. To explore the IMC proteome of the malaria parasite *Plasmodium falciparum* we applied a proximity-dependent biotin identification (BioID)-based proteomics approach, using the established IMC marker protein Photosensitized INA-Labelled protein 1 (PhIL1) as bait in asexual blood-stage parasites. Subsequent mass spectrometry-based peptide identification revealed enrichment of twelve known IMC proteins and several uncharacterized candidate proteins. We validated nine of these previously uncharacterized proteins by endogenous GFP-tagging. Six of these represent new IMC proteins, while three proteins have a distinct apical localization that most likely represent structures described as apical annuli in *Toxoplasma gondii*. Additionally, various Kelch13 interacting candidates were identified, suggesting an association of the Kelch13 compartment and the IMC in schizont and merozoite stages. This work extends the number of validated IMC proteins in the malaria parasite and reveals for the first time the existence of apical annuli proteins in *P. falciparum.* Additionally, it provides evidence for a spatial association between the Kelch13 compartment and the IMC in late blood-stage parasites.

## INTRODUCTION

*Plasmodium sp.* are members of the phylogenetic clade Alveolata that comprises a diverse group of unicellular eukaryotes including well-established phylogenetic groups such as Ciliates, Dinoflagellates and Apicomplexa (Cavalier-Smith, 1993). A defining feature of the Alveolata is a double-membrane system underlying the plasma membrane that is termed “alveoli” in ciliates and dinoflagellates (Allen, 1971; Lee and Kugrens, 1992) or collectively “inner membrane complex” (IMC) in apicomplexan parasites (Morrissette and Sibley, 2002). The IMC underlies the plasma membrane of the cell and consists of flattened double-membrane vesicles (Morrissette and Sibley, 2002) that are integrated into cytoskeletal components (Russell and Burns, 1984; Fowler *et al.*, 1998; Tran *et al.*, 2010). During evolution, this unique compartment was adapted to the individual ecological niches of the different clades (Kono *et al.*, 2013). This is reflected in the architecture and proteome of the IMC. For apicomplexans, the IMC has three main functions: i) it plays a major role in motility and invasion ii) it confers stability and shape to the cell, and iii) it provides a scaffolding framework during cytokinesis (Harding and Frischknecht, 2020; Ferreira *et al.*, 2021). While the IMC of the motile stages serves as the anchor for proteins involved in gliding motility and host cell invasion (Soldati *et al.*, 2004; Baum *et al.*, 2008; Perrin *et al.*, 2018), the gametocyte IMC appears to serve a structural role (Sinden, 1982; Dearnley *et al.*, 2012; Parkyn Schneider *et al.*, 2017). The IMC is generated *de novo* from Golgi-derived material in each replication cycle of the parasite during cytokinesis (Bannister *et al.*, 2000). This is similar to other compartments such as the secretory organelles that assemble a subset of parasite proteins crucial for invasion, egress and host cell modification (Blackman and Bannister, 2001).

To date about 45 IMC proteins have been identified in *Plasmodium* (Ferreira *et al.*, 2021), however, there is no comprehensive proteomic analysis of this structure available. From the confirmed IMC proteins it is evident that in addition to a common core set of conserved proteins found in all Alveolata, such as the Alveolins (Gould *et al.*, 2008; Gould *et al.*, 2011), the IMC includes many lineage-specific proteins, reflecting additional specialized roles (Kono *et al.*, 2012). A functional specialization is exemplified by a group of well-characterized proteins that form the so-called glideosome – the motor complex that drives the locomotion of all motile apicomplexan parasite stages (Gaskins *et al.*, 2004; Keeley and Soldati, 2004; Jones *et al.*, 2006) – anchored into the IMC. Another example of an apicomplexan-specific IMC protein is Photosensitized INA-Labelled protein 1 (PhIL1), which was first identified as an IMC protein in *Toxoplasma* (Gilk *et al.*, 2006) and is well characterized in both *Plasmodium* (Parkyn Schneider *et al.*, 2017; Saini *et al.*, 2017; Campelo Morillo *et al.*, 2020) and *Toxoplasma* (Gilk *et al.*, 2006; Barkhuff *et al.*, 2011). The *P. falciparum* PhIL1 (PF3D7_0109000) consists of 224 amino acids, was shown to be essential for mature gametocyte formation (Parkyn Schneider *et al.*, 2017) and was characterized as likely essential in blood-stage parasites (Zhang *et al.*, 2018). This is supported by the reported failed attempts in generating knockout parasites in *P. berghei* asexual blood stage(Saini *et al.*, 2017), even though no growth defect was observed upon glmS-based conditional knockdown in *P. falciparum* (Parkyn Schneider *et al.*, 2017). Proteins such as MORN1 (Gubbels *et al.*, 2006; Ferguson *et al.*, 2008), CINCH (Rudlaff *et al.*, 2019) or BTP1 (Kono *et al.*, 2016) define a subcompartment of the IMC termed the basal complex. The basal complex represents a dynamic ring structure and is implicated in organelle division of maturing daughter cells (Rudlaff *et al.*, 2019). The basal complex migrates distally away from the apical end of the daughter cells, marking the basal end of the newly formed parasite (Hu *et al.*, 2006; Gubbels *et al.*, 2006; Hu, 2008; Ferguson *et al.*, 2008; Kono *et al.*, 2016).

Additionally, another intriguing substructure is embedded in the apical area of the IMC– the apical annuli. These structures were first discovered in *T. gondii* (Hu *et al.*, 2006) and defined by cluster of rings with diameters ranging from 200 to 400 nm (Hu *et al.*, 2006; Engelberg *et al.*, 2019) containing *Tg*Centrin2 (Leung *et al.*, 2019; Lentini *et al.*, 2019) and at least six other apical annuli proteins (*Tg*AAP1-5, *Tg*AAMT) (Suvorova *et al.*, 2015; Engelberg *et al.*, 2019). AAPs, like many other IMC proteins, are specific for apicomplexan parasites and show some similarity to centrosomal proteins (Engelberg *et al.*, 2019). Two AAP homologues were identified in the genome of *P. falciparum* but have not been further characterized yet (Engelberg *et al.*, 2019).

Here we used proximity-dependent biotinylation with *Pf*PhIL1 to better characterize the IMC proteome and determine the interactome of this protein. This work identifies six novel IMC proteins, termed PhIL1 interacting candidates (PIC) as well as three putative apical annuli proteins, providing evidence for this structure in malaria parasites.

## RESULTS

### Identification of PhIL1-interacting proteins by proximity-dependent biotinylation (BioID)

The well-established, abundantly expressed *Pf*PhIL1 protein was used in order to identify putative PhIL1 interaction candidates and novel IMC proteins. For this, we employed proximity-dependent biotinylation (BioID (Roux *et al.*, 2012)) that exploits the activity of the biotin ligase BirA* for protein biotinylation and allows determination of putative protein-protein interaction by subsequent mass spectrometry.

First, we generated the transgenic parasite line *Pf*PhIL1-BirA-GFP by tagging the endogenous protein with BirA* and green fluorescent protein (GFP) using the selection-linked integration (SLI) system (Birnbaum *et al.*, 2017) (Figure S1A). Correct integration of the plasmid into the *Pf*PhIL1 gene locus was verified by PCR (Figure S1B). Expression of the fusion protein in schizont-stage parasites was confirmed by Western blot using anti-GFP antibodies (Figure S1C). Widefield fluorescence microscopy confirmed the expected IMC localization pattern (Parkyn Schneider *et al.*, 2017; Saini *et al.*, 2017; Campelo Morillo *et al.*, 2020) of PhIL1-BirA-GFP with an additional weak expression in ring stages (Figure 1A, S1D). Co-expression of the alveolin IMC protein ALV5 (Gould *et al.*, 2008; Hu *et al.*, 2010; Anderson-White *et al.*, 2011) in the PhIL1-BirA-GFP parasites revealed an IMC-typical localization of both tagged proteins with different fluorescence intensities at the parasite periphery of the developing merozoites (Figure S1E).

**Figure 1:**
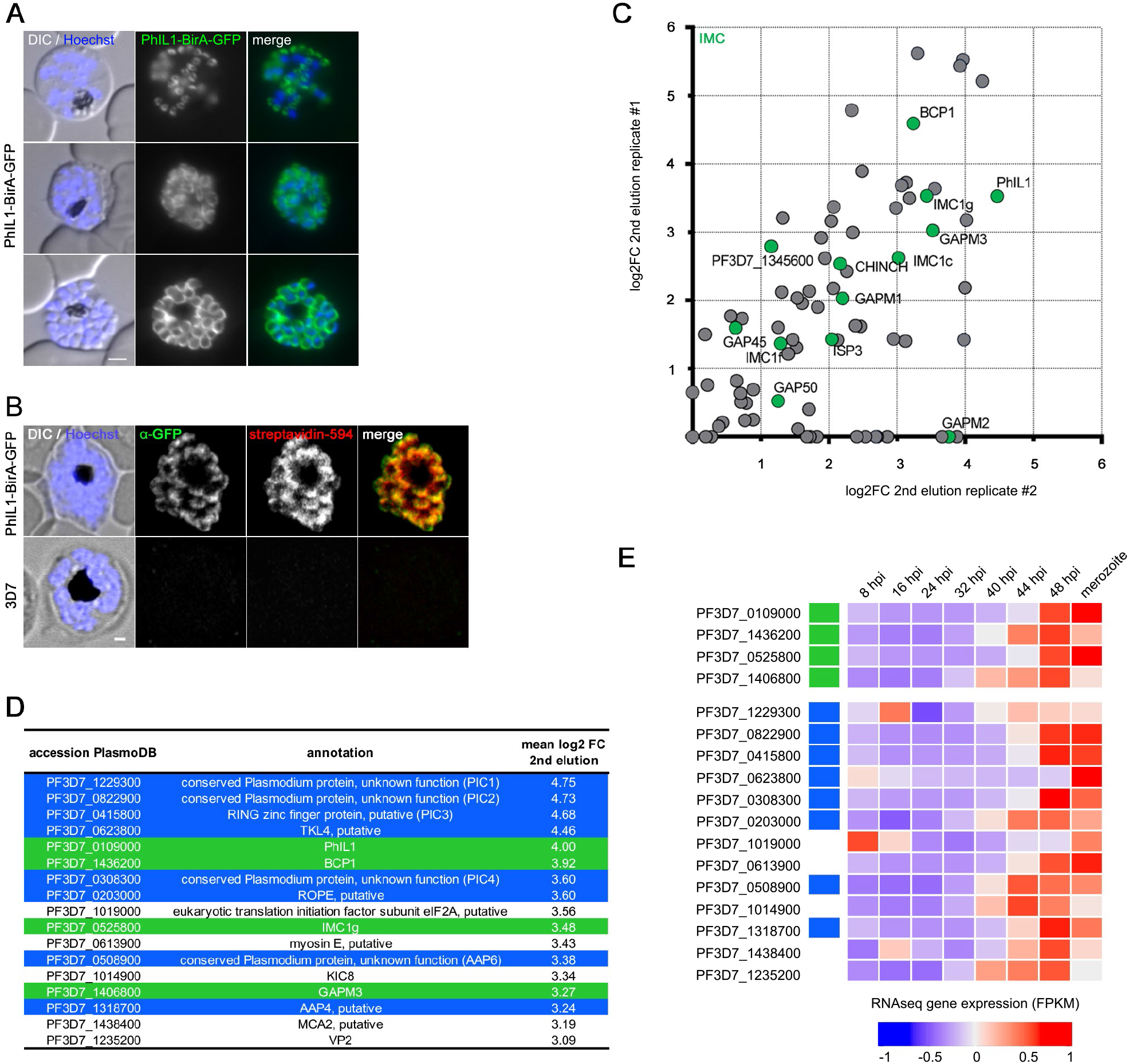
Identification of PhIL1 interacting protein using BioID. **(A)** IMC localisation of *Pf*PhIL1-BirA-GFP in schizonts and merozoites. Nuclei were stained with Hoechst-33342. Scale bar, 2 μm. **(B)** *Pf*PhIL1-based biotinylation at the IMC. *Pf*PhIL1-BirA-GFP (mouse anti-GFP, green) or 3D7 control parasites were fixed with MeOH at the schizont stage and biotinylated proteins were visualized by streptavidin-594 (red). Hoechst was used for DNA staining (blue). Scale bar, 1 μm. **(C)** Scatterplot showing proteins with a log2FC ≥ 0 in both BioID replicates. Green dots are indicating known IMC proteins. **(D)** Proteins with a mean log2FC> 3 in the *Pf*PhIL1-BirA-GFP compared to the 3D7 control sample are listed. For a complete list see Table S1. Known IMC proteins are highlighted in green, proteins marked in blue were selected for further validation. **(E)** Expression profiles(Wichers *et al.*, 2019) of the 17 most enriched proteins from (D) represented as a heat map (established IMC proteins (green), candidates selected for further validation (blue)).

Next, we induced proximity-dependent biotinylation by adding biotin to tightly synchronized *Pf*PhIL1-BirA-GFP and 3D7 control parasite cultures 40 hours post invasion (hpi). Using fluorescently labelled streptavidin, we detected *Pf*PhIL1-based biotinylation at the IMC in the presence of biotin in the transgenic parasite line, while no biotin signal was visible in 3D7 control parasites (Figure 1B). Subsequently, biotinylated proteins were affinity-purified from schizont-stage parasite lysate and subjected to mass-spectrometry based protein identification resulting in the identification of 219 proteins with a mean log_2_ fold change >1 in the first elution and 58 proteins in the second elution. Proteins found to be enriched compared to the 3D7 control are shown in Figure 1C,D, Figure S1F and Table S1. The top 17 enriched proteins show a mean log_2_ fold change> 3 and include four proteins that have been previously localized to the IMC or the basal complex: BCP1 (Rudlaff *et al.*, 2019), ALV3 (Gould *et al.*, 2008; Hu *et al.*, 2010; Anderson-White *et al.*, 2011), GAPM3 (Bullen *et al.*, 2009) and the bait protein PhIL1 (Parkyn Schneider *et al.*, 2017; Saini *et al.*, 2017) (Figure 1D).

Genes coding for IMC proteins are typically characterized by a uniform transcriptional profile during asexual replication cycle with a maximum expression in the late schizont/merozoite stage (Hu *et al.*, 2010). To filter out potential IMC proteins we thus discriminated enriched proteins by comparative analysis of the expression pattern of all candidates (Figure 1E). All genes except PF3D7_1019000 (putative eukaryotic translation initiation factor subunit eIF2A) exhibit the expected profile, therefore PF3D7_1019000 was excluded from further analysis.

Additionally, the following proteins were excluded based on published literature: PF3D7_1235200 (VP2) was previously localized to the parasite plasma membrane or punctate intracellular inclusions (McIntosh *et al.*, 2001), PF3D7_1014900 (KIC8) and PF3D7_1438400 (MCA2) are linked with the Kelch13 compartment(Birnbaum *et al.*, 2020) and PF3D7_0613900 (MyoE) was shown to have a non-IMC like localization in *P. berghei* schizonts (Wall *et al.*, 2019), resulting in a list of eight putative novel IMC proteins (PF3D7_1229300, PF3D7_0822900, PF3D7_0415800, PF3D7_0623800, PF3D7_0308300, PF3D7_0203000, PF3D7_0508900, PF3D7_1318700). Additionally, we included PF3D7_1310700 and PF3D7_0530300, which were – together with the known IMC protein GAPM2 (Bullen *et al.*, 2009) – highly enriched in replicate #2 (log2 fold change of 2.86 and 2.69), but absent in the eluate of replicate #1 (Table S1). This resulted in a final selection of ten candidates for further analysis.

### Localization of PhIL1-interacting candidates (PIC) reveals six novel IMC proteins

Based on their potential interaction with PhIL1, we named the previously non-annotated proteins PhIL1-interacting candidates (PIC). The ten selected candidate proteins were further validated. We first aimed to localize them by endogenous C-terminal GFP tagging using the SLI system. We achieved successful modification of the respective gene loci for eight candidate proteins (Figures 2, 3, 4), while two genes, *Pfrope* (Werner *et al.*, 1998) (PF3D7_0203000) and *pic5* (PF1310700), were refractory to 3’modifications using the SLI system. From the resulting eight endogenously tagged cell lines, seven expressed the fusion protein to a sufficient level that allowed its subcellular localization, even though PIC3-GFP and PIC6-GFP expression levels were very weak (Figure 2 A–E, Figure S2). The TKL4 (Abdi *et al.*, 2010)-GFP (PF3D7_0623800) expression level was too low for conclusive localization (Figure S2D). Fluorescence microscopy of PIC1-4 and 6 showed an IMC-typical localization pattern (Figure 3A-E, S2). We verified the IMC localization by co-expression of the IMC marker protein ALV5 in the GFP-tagging cell lines (Figure 3).

**Figure 2:**
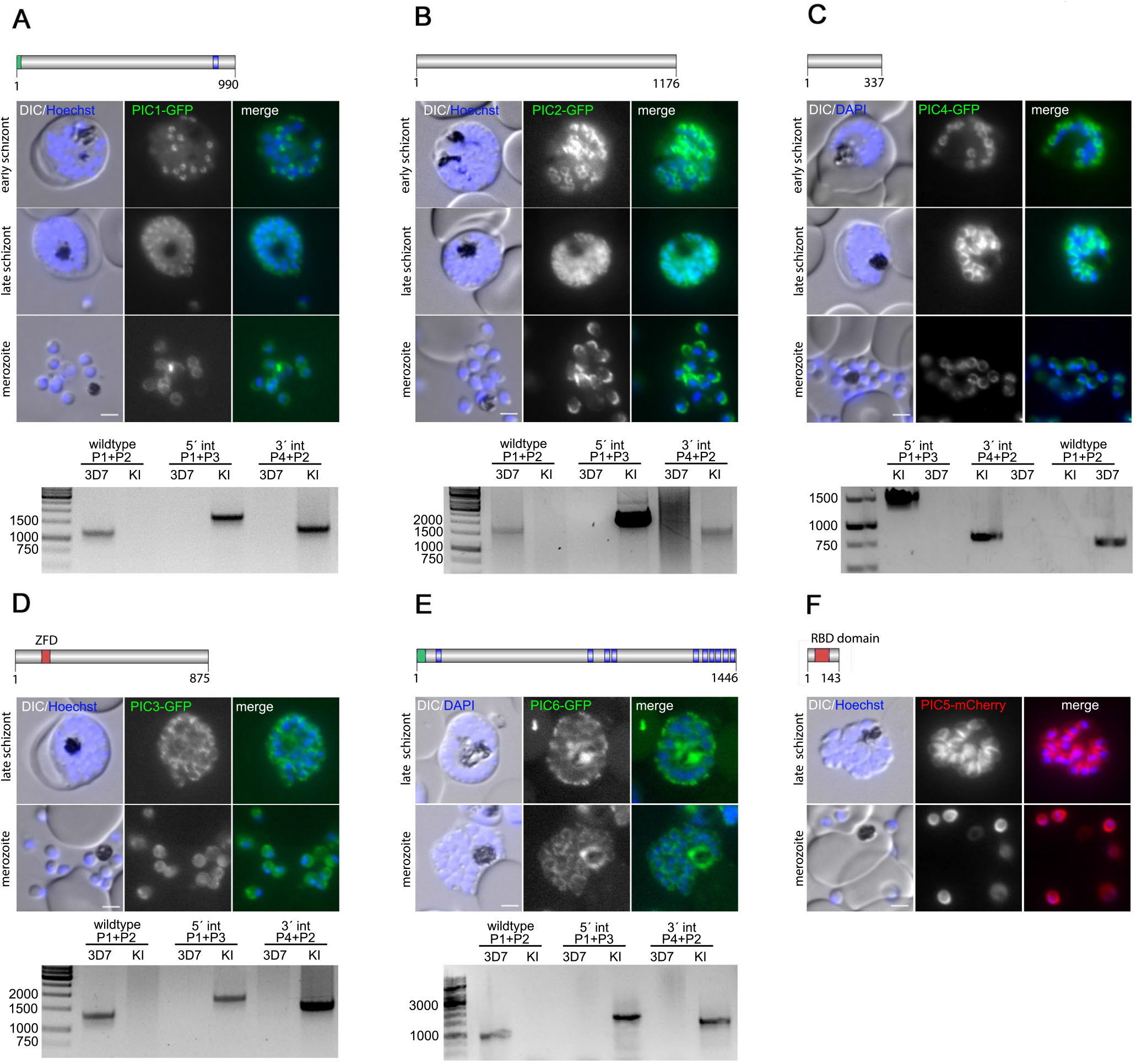
Localization of PhIL1 interacting candidates (PIC) using endogenous GFP tagging. **(A)** PIC1 (PF3D7_1229300), **(B)** PIC2 (PF3D7_0822900), **(C)** PIC4 (PF3D7_0308300), **(D)** PIC6 (PF3D7_0530300), **(E)** PIC3 (PF3D7_0415800): Schematic representation of the protein (protein length indicated as number of amino acids) with putative protein domains (blue, transmembrane domain; green, signal peptide; red, Zinc Finger domain (ZFD) or RBD, RNA binding domain). Localization of PIC-GFP fusion proteins in schizonts and free merozoites, PCR-based confirmation of the correct insertion of GFP-encoding integration plasmid into the targeted loci. Ladder size indicated in base pairs (bp). gDNA from parental 3D7 was used as control. KI = knock-in **(F)** Localization of PIC5-(PF3D7_1310700)-mCherry fusion proteins episomally expressed under the control of the late-stage promoter *ama1* in schizonts and free merozoites. Nuclei stained with Hoechst-33342 or DAPI. Zoom factor: 400%. Scale bar 2 μm.

**Figure 3:**
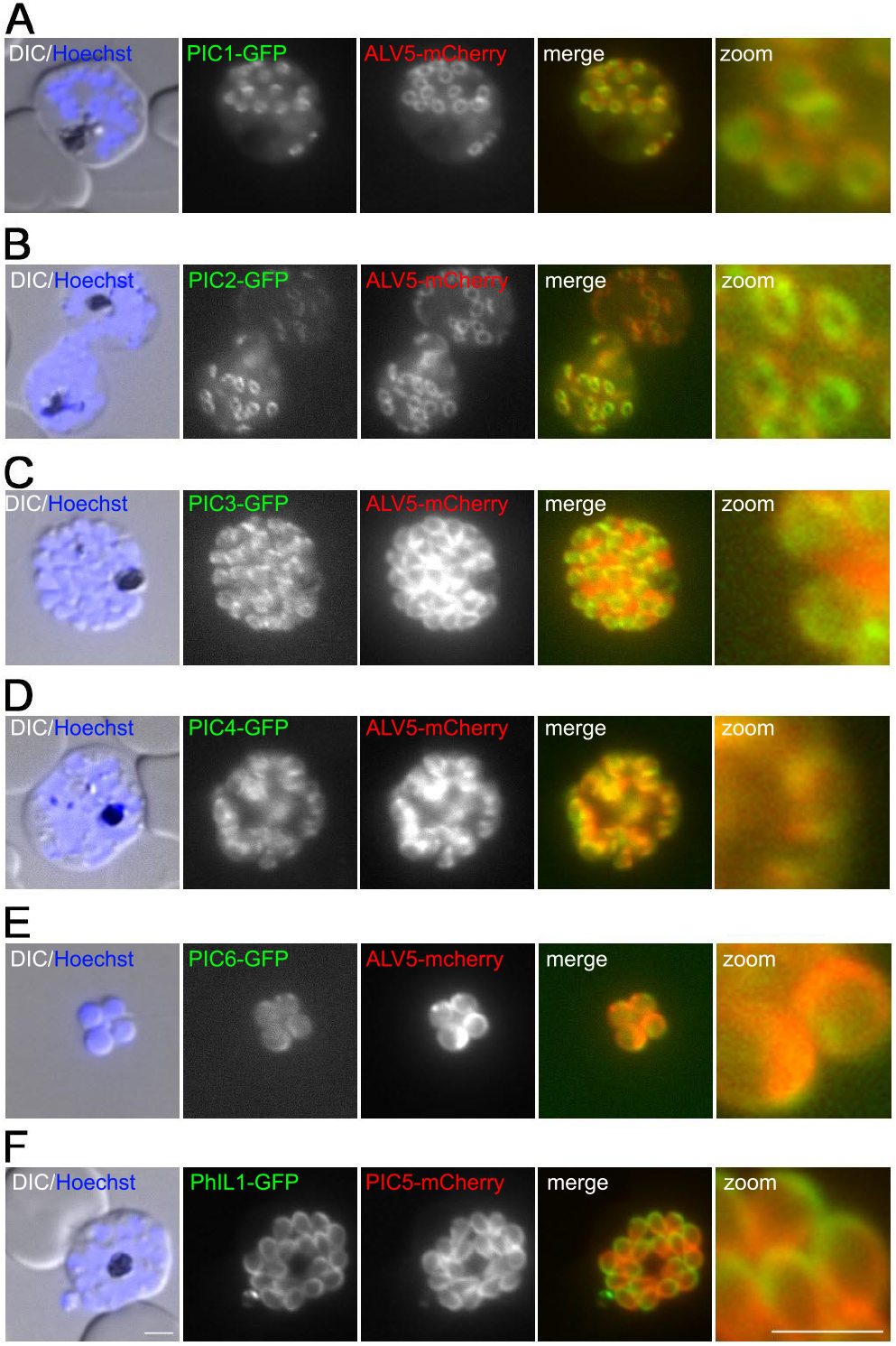
Co-expression of PICs with the IMC marker ALV5-mCherry Co-**expression** of the candidate proteins PIC1. **(A)**, PIC2 **(B)**, PIC3 **(C)**, PIC4 **(D)**, PIC6 **(E)**, and the IMC marker protein ALV5 episomally expressed as mCherry fusion protein under the control of the late-stage promoter *ama1*. (F) PIC5 episomally expressed as mCherry fusion protein under the control of the late-stage promoter *ama1* in PhIL1-BirA-GFP parasites. Nuclei stained with Hoechst-33342. Zoom factor: 400%. Scale bars 2 μm.

**Figure 4:**
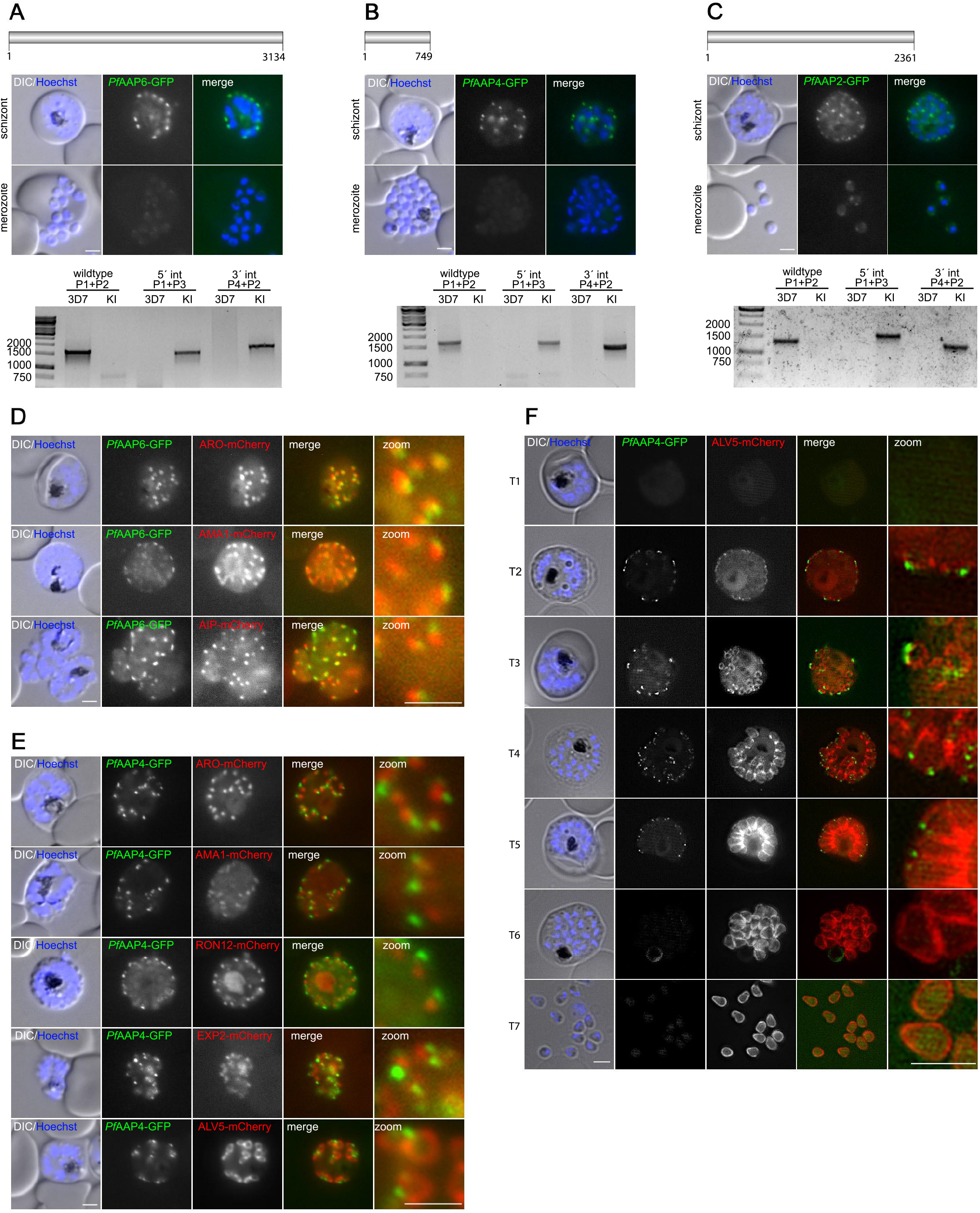
*Pf*AAP6, *Pf*AAP4 and *Pf* AAP2 define a novel, apical structure in *P. falciparum*. **(A)** *Pf*AAP6 (PF3D7_0508900), **(B)** *Pf*AAP4 (PF3D7_1318700), **(C)** *Pf*AAP2 (PF3D7_1312800): Schematic representation of each protein (protein length indicated as number of amino acids). Localization of AAP-GFP fusion proteins in schizonts and free merozoites, PCR-based confirmation of the correct insertion of GFP-encoding integration plasmid into the targeted loci. Ladder size indicated in base pairs (bp) gDNA from parental 3D7 was used as control. Nuclei stained with Hoechst-33342. Scale bars 2 μm. **(D)** Co-localization of *Pf*AAP6-GFP with the rhoptry bulb marker protein ARO-mCherry, the microneme marker AMA1-mCherry and the rhoptry neck marker AIP-mCherry. **(E)** Co-localization of *Pf*AAP4-GFP with the rhoptry bulb marker protein ARO-mCherry, the microneme marker AMA1-mCherry, the rhoptry neck marker RON12-mCherry, the dense granules marker EXP2-mCherry and the IMC marker ALV5-mCherry. **(F)** Structured illumination microscopy (SIM) images of *Pf*AAP4-GFP with the IMC marker ALV5-mCherry. T1-T3 showing early schizonts, T4-T6 late schizonts and T7 free merozoites. Zoom factor: 400%. Scale bars 2 μm.

Additionally, due to the lack of SLI-based integration into the PIC5 (PF3D7_1310700) locus, this protein was episomally expressed as an mCherry fusion in the *Pf*PhIL1-BirA-GFP cell line. Again, an IMC-typical localization pattern was observed (Figure 2F, 3) resulting in a total of six PIC proteins with IMC localization (PIC1-6).

### Distinct localization of apical annuli protein homologues in *P. falciparum*

Two other PICs identified by PhIL1-based BioID, PF3D7_0508900-GFP and PF3D7_1318700-GFP (annotated in PlasmoDB as *Pf*AAP4) showed a unique localization pattern clearly distinct from their IMC-localized counterparts (Figure 4A–B, S3).

*Pf*AAP4 is the homologue of the apical annuli protein *Tg*AAP4 (TGGT1_230340) that was recently described in *T. gondii* (Engelberg *et al.*, 2019; Barylyuk *et al.*, 2020). Homology searches of the list of enriched proteins from the BioID experiment for further homologues of *T. gondii* apical annuli proteins retrieved PF3D7_1312800 (*Tg*AAP2, TGGT1_295850) and PF3D7_1455200 *(Tg*AAMT, TGGT1_310070) (Figure S4). Accordingly, we named these two proteins *Pf*AAP2 and *Pf*AAMT. PF3D7_0508900 was annotated as *Pf*AAP6, due to its localization similar to *Pf*AAP2 and *Pf*AAP4. Next, we generated an endogenously tagged *Pf*AAP2-GFP cell line (Figure 4C), which revealed a similar localization to *Pf*AAP6-GFP and *Pf*AAP4-GFP. These three GFP fusion proteins localize at the apical pole in nascent daughter cells (Figure 4 A–C, S3). While *Pf*AAP4-GFP could not be visualized in free merozoites (Figure 4B), *Pf*AAP6-GFP and *Pf*AAP2-GFP showed a weak signal (Figure 4A,C, S3). In order to exclude a localization with the apical organelles we co-localized *Pf*AAP6-GFP and *Pf*AAP4-GFP with appropriate marker proteins: *Pf*ARO (Cabrera *et al.*, 2012; Geiger *et al.*, 2020) (rhoptry bulb marker), *Pf*RON12 (Knuepfer *et al.*, 2014; Ito *et al.*, 2019) or *Pf*AIP (Geiger *et al.*, 2020) (rhoptry neck marker) and *Pf*AMA1 (Peterson *et al.*, 1989; Healer *et al.*, 2002) (micronemes marker) and additionally *Pf*AAP4-GFP with *Pf*EXP2 (Bullen *et al.*, 2012; Mesén-Ramírez *et al.*, 2019) (dense granules marker). To achieve this, we transfected the endogenously tagged *Pf*AAP6-GFP and *Pf*AAP4-GFP expressing cell lines with an expression plasmid for the respective mCherry fusion proteins (Figure 4D-E). This revealed that localization of both *Pf*AAP6-GFP and *Pf*AAP4-GFP is clearly distinct from – but spatially associated with – the apical organelles.

Super-resolution microscopy indicated that the apical annuli structures are embedded in the IMC in *T. gondii* (Engelberg *et al.*, 2019). To probe into the spatial relation of the *Pf*AAP4 defined apical annuli structure and the IMC in *P. falciparum*, we co-expressed it with the IMC marker protein ALV5 using fluorescence (Figure 4E) and SIM super-resolution microscopy. IMC and apical annuli develop at the same time during early schizont development at the apical pole of nascent daughter cells. While the punctuated *Pf*AAP4-GFP signal remains restricted to the apical area the ALV5-mCherry signal expands with the IMC from the apical to the basal pole during the maturation of the daughter cells during late schizogony. While the *Pf*AAP4-GFP signal is not in all instances clearly assigned to the ALV5-mCherry defined IMC structure, the putative annuli are clearly in vicinity of the IMC and part of the expanding circular structure of the IMC (see T3 and T4 in Figure 4F). This suggests an embedment or the association of the *Pf*AAP4 structure with the IMC. In late stage schizonts and free merozoites the *Pf*AAP4 signal is lost (Figure 4F).

### Functional assessment of PIC1, PIC2, PIC3, PIC4, *Pf*AAP4, *Pf*AAP6 and PhIL1

The introduction of a glmS ribozyme (Prommana *et al.*, 2013) sequence upstream of the 3’ untranslated region allowed the conditional degradation of mRNA of PIC1, PIC2, PIC3, *Pf*AAP4, and *Pf*AAP6 upon glucosamine addition. First, we quantified the reduction of the GFP signal in the presence of 2.5 mM glucosamine which showed that this leads to an 85.33% (SD ±1.97) reduction of GFP fluorescence intensity in PIC1-GFP and to 46.39% (SD ±0.57) reduction in PIC3-GFP (Figure S5A,B), but no reduction in PIC2-GFP and *Pf*AAP4-GFP. The expression of *Pf*AAP6-GFP in the absence of glucosamine was already too low to be reliably quantified (Figure S5D–F). Subsequently, for those cell lines that showed a significant reduction in the expression of endogenously tagged protein, PIC1-GFP and PIC3-GFP, parasite proliferation was analyzed in absence and presence of glucosamine. No growth defect could be observed (Figure S5A,B).

Due to the candidate-dependent knock-down efficacy with the glmS-based knock-down approach we also applied a conditional inactivation via the knockside-ways(Birnbaum *et al.*, 2017) approach for one IMC protein (PIC4). In contrast to the RNA-based glmS method, this system allows the mislocalization of a protein from its site of action into the nucleus upon addition of rapalog and was shown highly effective for membrane-associated proteins such as *Pf*Rab5a (Birnbaum *et al.*, 2017), *Pf*KELCH13 (Birnbaum *et al.*, 2020) and *Pf*AIP (Geiger *et al.*, 2020). To achieve mislocalization, we used our PIC4-2xFKBP-GFP-2xFKBP parasites line and transfected it with the 1xNLS-FRB-mCherry plasmid (Birnbaum *et al.*, 2017) encoding the nuclear mislocalizer coupled to FRB. Efficient mislocalization of PIC4-2xFKBP-GFP-2xFKBP into the nucleus upon addition of rapalog was shown by live-cell epifluorescence microscopy, but this did not translate into any decreased parasite proliferation (Figure S5C). Taken together, functional inactivation of PIC1, PIC3 or PIC4 by either ribozyme mediated RNA degradation or knock sideways had no measurable growth phenotype and argues for the redundancy of these IMC proteins in asexual proliferation. This was further supported by the successful disruption of PIC4 by generating a cell line expressing a truncated version of the endogenous PIC4 (PIC4-TGD) using the SLI (Birnbaum *et al.*, 2017) system (Figure S6A–C). Additionally, we also assessed the essentiality of *Pf*PhIL1 for asexual development of the parasite using SLI-TGD (Figure S6D–F). The growth rate of *Pf*PhIL1-TGD parasites was reduced by 29% over two cycles compared to wild-type parasites (Figure S6D).

### The Kelch13 compartment is IMC associated in merozoites

Artemisinin resistance in *P. falciparum* is mainly associated with mutations in the parasite *pfkelch13* gene(Ariey *et al.*, 2014; Straimer *et al.*, 2015). Kelch13 has been shown to be involved in endocytosis at the ring stage (Birnbaum *et al.*, 2020) and to be essential for parasite growth (Birnbaum *et al.*, 2017). Endogenously tagged Kelch13 was originally localized to foci in close proximity to the parasite’s food vacuole and showed multiple foci in schizonts (Birnbaum *et al.*, 2017). Subsequent work also revealed an association of Kelch13 with **~**170 nm diameter doughnut-shaped structures at the parasite periphery (Yang *et al.*, 2019) and no association with known markers of the secretory system (Birnbaum *et al.*, 2020; Gnädig *et al.*, 2020).

KIC8 (Kelch13 interaction candidate 8), KIC2 (Kelch13 interaction candidate 2) (Birnbaum *et al.*, 2020) and MCA2 (metacaspase-2) are among the top 25 enriched proteins in our PhIL1-based BioID assay. Vice versa, KIC8, KIC2, MCA2, as well as PhIL1 were found to be enriched in a Kelch13 based DiQ-BioID experiment (Birnbaum *et al.*, 2020). KIC8 and KIC2 were shown to co-localize with Kelch13 (Birnbaum *et al.*, 2020). To investigate a link between the IMC and the Kelch13 compartment we analyzed the list of enriched proteins in our PhIL1-based BioID approach for other members of the Kelch13 compartment. Indeed, almost all KIC proteins are significantly enriched (Figure 5A,B). In the first elution Kelch13 and all KICs except UBP1 and KIC10 were enriched. In line with this, KIC10 was previously identified as not co-localizing with the Kelch13 compartment in schizonts (Birnbaum *et al.*, 2020).

**Figure 5:**
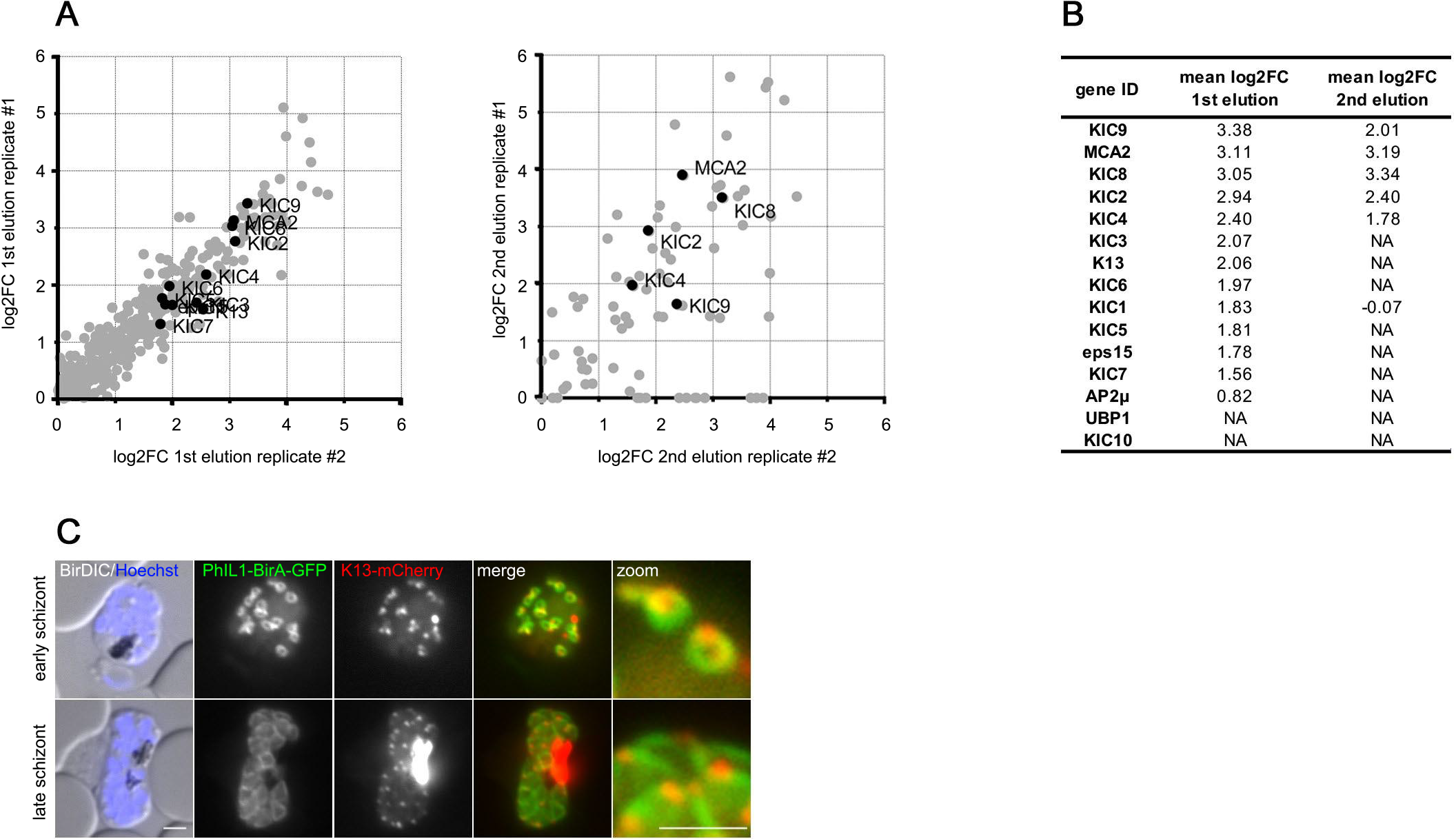
The Kelch13 compartment is IMC associated. **(A)** Scatterplot showing proteins with a log2FC ≥ 0 in both BioID replicates (first and second elution). Kelch13 interacting candidates(Birnbaum *et al.*, 2020) are highlighted. **(B)** Mean log2FC values for all KIC proteins in both BioID fractions **(C)** Localization of Kelch13-mCherry episomally expressed under the control of the *nmd3* promoter in the PhIL1-BirA-GFP cell line. Nuclei stained with Hoechst-33342. Zoom factor: 400%. Scale bars 2 μm. NA = not applicable

To further probe into this intriguing localization, we first aimed to co-localize KIC8 with PhIL1 by dual tagging of these proteins, unfortunately without success. We therefore overexpressed mCherry-Kelch13, a construct known to co-localise with endogenous Kelch13 (Birnbaum *et al.*, 2020) in the PhIL1-BirA-GFP expressing cell line to enable co-localization. As shown in Figure 5C Kelch13 is intimately associated with the nascent IMC at the apical pole during daughter cell formation and in mature schizonts supporting putative PhIL1 interaction This indicates that either the Kelch13 compartment is intimately associated with the IMC or that Kelch13 compartment proteins are involved in other IMC related functions in late blood-stage parasites.

## DISCUSSION

In the present study we used BioID to probe into the proteome of the IMC in late blood stages of *P. falciparum*. Within the top 58 enriched proteins that show a mean log_2_ fold change > 1 are twelve known IMC proteins (PhIL1 (Parkyn Schneider *et al.*, 2017; Saini *et al.*, 2017), CHINCH (Rudlaff *et al.*, 2019), GAPM1-3 (Bullen *et al.*, 2009; Kono *et al.*, 2012), ALV4/IMC1g (Hu *et al.*, 2010), ALV5/IMC1c (Hu *et al.*, 2010), IMC1f (Al-Khattaf *et al.*, 2015), GAP45 (Baum *et al.*, 2006), PF3D7_1345600 (Kono *et al.*, 2012), BCP1 (Rudlaff *et al.*, 2019), ISP3 (Kono *et al.*, 2012)) comprising approximately one quarter of all known IMC proteins (Ferreira *et al.*, 2021). Additionally, the BioID-based protein identification revealed two proteins, PPP8 (Rudlaff *et al.*, 2019) and PF3D7_1327300 (Parkyn Schneider *et al.*, 2017), that were previously identified using immunoprecipitation (IP) approaches. Of note PIP1-3 (PhIL1 interacting proteins 1-3) identified in gametocytes (Parkyn Schneider *et al.*, 2017) were not among the hits, which might indicate different IMC composition in schizonts and gametocytes.

We validated 9 candidates identified by our BioID approach and present six novel *P. falciparum* IMC proteins we named PIC 1-6 (**P**hiL1 **I**nteracting **C**andidates). PIC2 (PF3D7_0822900) has been previously identified in IP experiments using *Pf*MyoA (Green *et al.*, 2017) or *Pf*PhIL1 (Parkyn Schneider *et al.*, 2017) / *Pb*PhIL1 (Saini *et al.*, 2017) as bait, while PIC5 (PF3D7_1310700) – which contains a predicted putative RNA binding domain (Mitchell *et al.*, 2019) – was present in IP experiments of *Pb*PhIL1 (Saini *et al.*, 2017) and *Pf*CINCH (Rudlaff *et al.*, 2019).

PIC1 and PIC6 contain a predicted (Almagro Armenteros *et al.*, 2019) N-terminal signal peptide (Figure 2A and E). This N-terminal signature is rather unusual for IMC proteins and only the known IMC proteins NIPA (PF3D7_0522600) and DHHC9 (PF3D7_1115900) also show this feature (Almagro Armenteros *et al.*, 2019). With PIC1, PIC3 and PIC6 three proteins contain at least one predicted (Sonnhammer *et al.*, 1998) transmembrane domain (Figure 2 A,D,E). Single putative transmembrane domains are known for the IMC protein G2 (PF3D7_0929600), while multiple transmembrane domains are also predicted for GAPM1-3, DHHC1-3 and DHHC9, BTP1/BTP2, NIPA and GAP40/GAP50.

Functional inactivation of PIC1, PIC3 or PIC4 by either ribozyme-mediated mRNA degradation or knock sideways revealed no growth phenotype. While PIC1 and PIC3 have been classified as dispensable in a genome-wide mutagenesis screen(Zhang *et al.*, 2018), PIC4 is predicted to be essential. Although ribozyme mediated protein down-regulation was substantial and mis-localization was efficient (Figure S5), the residual amount of these proteins still present might be sufficient to mediated normal IMC function under the given growth conditions. However, as the gene encoding PIC4 could be disrupted, this is not a likely explanation for this candidate and it can be assumed that PIC4 is dispensable.

We also probed into the essentiality of *Pf*PhIL1 itself and the successful generation of a *Pf*PhIL1 deficient parasite line reveals that *Pf*PhIL1 is not essential for *in-vitro* growth of asexual parasites either. However, the *Pf*PhIL1 knockout parasites showed a reduced growth rate indicating a fitness loss of this cell line. This finding is in agreement with observed reduced fitness upon knockout of *Tg*PhIL1 in *T. gondii* (Barkhuff *et al.*, 2011).

Beside novel IMC proteins the BioID list contains at least three putative apical annuli proteins of *P. falciparum (Pf*AAP2*, Pf*AAP4*, Pf*AAP6), for which endogenous GFP-tagging revealed an apical localization but no co-localization with the apical organelles. Apical annuli or peripheral annuli are structures that so far have only been described in *T. gondii* and were discovered by localization of *Tg*CEN2 (Hu *et al.*, 2006) and *Tg*AAP1 (Suvorova *et al.*, 2015). They consist of five to six rings at the lower edge of the apical cap. While the function of the apical annuli remains unresolved they have been linked with the IMC, more precisely with the *T. gondii* ISP subcompartment (Beck *et al.*, 2010). Several IMC proteins (including *Tg*PhIL1) were enriched in a *Tg*AAP4 BioID (Engelberg *et al.*, 2019) and *Tg*AAP4 has been previously identified in a BioID experiment using the alveolar suture component *Tg*ISC4 (Chen *et al.*, 2017) as bait. Although their function is unknown it has been speculated that apical annuli may serve a pore function facilitating efficient exchange of nutrients and waste products across the IMC or may serve as a signalling platform (Engelberg *et al.*, 2019). So far eight proteins of *T. gondii* have been shown to localize to the annuli (*Tg*AAP1-5 (Engelberg *et al.*, 2019), *Tg*AAMT (Engelberg *et al.*, 2019), *Tg*CEN2 (Hu *et al.*, 2006; Leung *et al.*, 2019; Lentini *et al.*, 2019), *Tg*PI-PLC (Hortua Triana *et al.*, 2018)).

The absence of three of five *aap* genes - present in *T. gondii* (Engelberg *et al.*, 2019) *-* in the *Plasmodium* genome and the lack of a homologue of *Pf*AAP6 in *T. gondii* raises the question how conserved the proteome and the function of the annuli is between these species. Further studies are needed to address these questions.

Of note apical annuli proteins are expressed in multiple stages of the *P. falciparum* life cycle, as indicated by high mRNA expression of *pfaap4* in schizonts, merozoites (Wichers *et al.*, 2019), gametocytes (López-Barragán *et al.*, 2011), ookinetes (López-Barragán *et al.*, 2011) and sporozoites (Zanghì *et al.*, 2018) even though we did not observe a detectable signal of *Pf*AAP4–GFP, *Pf*AAP6-GFP or *Pf*AAP2-GFP in gametocytes (Figure S3).

Additionally, we identified *Pf*AAMT (PF3D7_1455200) as a homologue of *Tg*AAMT (Engelberg *et al.*, 2019), indicating that the list might contain additional *P. falciparum* apical annuli proteins. Together with recent reports on conserved proteins of the conoid structure (Wall *et al.*, 2016; Leung *et al.*, 2020; Koreny *et al.*, 2020; Bertiaux *et al.*, 2020) the presence of apical annuli in *P. falciparum* may indicate a similar and conserved organization of the apical pole in Apicomplexan parasites.

An additional, interesting finding of our PhIL1-based BioID assay was the significant enrichment of Kelch13 interacting candidates (KICs). This is in agreement with the Kelch13 based BioID approach (Birnbaum *et al.*, 2020), which identified PhIL1, as well as other IMC proteins such as ALV4/IMC1g, ALV5/IMC1c, PF3D7_1345600, ISP3 and GAPM2 to be enriched although at lower level than other KICs. We show intimate association between the Kelch13 compartment with the nascent IMC at the apical pole during daughter cell formation (Figure 5C). Kelch13 has been recently suggested to localize to cytostome-like structures at the parasite periphery in schizonts (Yang *et al.*, 2019). The function of the Kelch13 compartment and cytostome-like structures in late schizont and merozoites remains to be elucidated. It could either represent a re-purposing of these endocytosis-linked structures (Spielmann *et al.*, 2020) for host cell invasion and the establishment of the parasite in a new host cell or serve an endocytosis-related function in schizonts, as it has been recently shown that schizonts still appear to endocytose host cell hemoglobin (Bisio *et al.*, 2020).

## Authors’ contributions

Conceptualization: TWG; Methodology: JSW, JW, DH, SP, JS, PB, TS, CL, MF, AB, TWG; Validation: JSW, JW, DH, SP, JS, MS, JK, LW, SS, HvT, PB; Formal Analysis: JSW, JW, DH, SP, JS, TWG; Writing JSW, JW, TWG; Visualization: JSW, JW, JS, TWG; Funding Acquisition: JS, CL, MF, AB, TWG; Resources: CL, TWG; Project Administration: TS, CL, MF, AB, TWG; Supervision: TS, CL, AB TWG. All authors read and approved the manuscript.

## Acknowledgements

The authors thank Michael Geiger for cloning of pARL-ama1_AIPmCherry, Jacobus Pharmaceuticals for WR99210, the Proteomics Core Facility at EMBL Heidelberg for support with mass spectrometry sample preparation, measurements and data analysis. Microscopy experiments were supported by the Advanced Light and Fluorescence Microscopy facility at the CSSB, in particular Roland Thünauer. The following reagent was obtained through BEI Resources, NIAID, NIH: DSM1, MRA-1161.

## METHODS

### *P. falciparum* culture

The *Plasmodium falciparum* clone 3D7 (Walliker *et al.*, 1987) was cultured at a hematocrit of 5% in human O^+^ erythrocytes according to standard procedures^6^. To maintain synchronized parasites, cultures were treated with 5% sorbitol (Lambros and Vanderberg, 1979). The PKG-inhibitor compound 2 was used at a final concentration of 1 μM in order to arrest parasite egress before exoneme/microneme secretion as previously described (Collins *et al.*, 2013).

### Cloning of DNA constructs

For generating the PhIL1-BirA-GFP construct, a homology region of 669 bp was amplified using 3D7 gDNA and cloned into pSLI-TGD (Birnbaum *et al.*, 2017) via NotI and AvrII restriction sites, next the BirA sequence was amplified from pSP-GFP-BirA (Khosh-Naucke *et al.*, 2018) and inserted via the AvrII/MluI restriction site, resulting in pSLI-PhIL1-BirA-GFP. For endogenous tagging using the SLI system (Birnbaum *et al.*, 2017), a homology region of 426 – 1087 bp (1001 bp for *Pf*3D7_1229300, 1087 bp for *Pf*3D7_0822900, 1056 bp for *Pf*3D7_0415800, 866 bp for *Pf*3D7_0623800, 1028 bp for *Pf*3D7_0203000, 1041 bp for *Pf*3D7_0508900, 917 bp for *Pf*3D7_1318700, 426 bp for *Pf*3D7_1310700 and 1038 bp for *Pf*3D7_1312800 respectively) was amplified using 3D7 gDNA and cloned into pSLI-GFP-glmS (Burda *et al.*, 2020) (derived from pSLI-GFP (Birnbaum *et al.*, 2017)) using the NotI/MluI restriction site. For PF3D7_0308300, a 650 bp homology region was amplified from 3D7 gDNA and cloned into pSLI-sandwich (Birnbaum *et al.*, 2017) via NotI and AvrII restriction sites. For targeted gene disruption (TGD) a homology region of 348 bp for *Pf*PhIL1 or 273 bp for PF3D7_0308300 were amplified from 3D7 gDNA or synthesized by Life Technologies (Darmstadt, Germany) and cloned into pSLI-TGD using the NotI/MluI restriction site (Birnbaum *et al.*, 2017).

For co-localization experiments, the full length sequence of organellar marker genes for micronemes (*ama1 Pf*3D7_1133400) (Peterson *et al.*, 1989; Healer *et al.*, 2002), rhoptry neck (*aip*, *Pf*3D7_1136700; *ron12 Pf*3D7_1017100) (Knuepfer *et al.*, 2014; Ito *et al.*, 2019; Geiger *et al.*, 2020), inner membrane complex (*imc1c/alv5, Pf*3D7_1003600) (Hu *et al.*, 2010) were amplified from 3D7 cDNA and for dense granules (*exp2,* PF3D7_1471100) (Bullen *et al.*, 2012) from the plasmid pEXP2rec (Mesén-Ramírez *et al.*, 2019) and cloned into pARL-ama1-ARO-mCherry-BSD (Cabrera *et al.*, 2012) using the AvrII/KpnI restriction site or in pARL-ama1-ARO-mCherry-yDHODH (derived from pARL-ama1-ARO-mCherry-BSD (Cabrera *et al.*, 2012)).

Additionally, the plasmids pBcamR-alv5-mCherry (Kono *et al.*, 2012), pARL-ARO-mCherry-BSD (Cabrera *et al.*, 2012) were used for co-localization experiments. Oligonucleotides used for cloning are listed in Table S2.

### Transfection of *P. falciparum*

For transfection, Percoll-purified (Rivadeneira *et al.*, 1983) parasites at late schizont stage were transfected with 50 μg plasmid DNA using Amaxa Nucleofector 2b (Lonza, Switzerland) as previously described (Moon *et al.*, 2013). Transfectants were selected using either 4 nM WR99210 (Jacobus Pharmaceuticals), 0.9 μM DSM1 (BEI Resources) or 2 μg/mL blasticidin S (Life Technologies, USA). In order to select for parasites carrying the genomic modification via the SLI system(Birnbaum *et al.*, 2017), G418 (ThermoFisher, USA) at a final concentration of 400 μg/mL was added to a culture with 5% parasitemia. The selection process and integration test were performed as previously described (Birnbaum *et al.*, 2017).

### Imaging and immunofluorescence assay (IFA)

Fluorescence images were captured using a Zeiss Axioskop 2plus microscope with a Hamamatsu Digital camera (Model C4742-95). Microscopy of live parasite-infected erythrocytes was performed as previously described(Grüring and Spielmann, 2012). Briefly, parasites were incubated in standard culture medium with 1 μg/mL Hoechst-33342 (Invitrogen) for 15 minutes at 37°C prior to imaging. 5.4 μL of infected erythrocytes were added on a glass slide and covered with a cover slip. IFAs were performed as described previously(Bachmann *et al.*, 2015). Mouse anti-GFP antibodies (Roche) were used at a dilution of 1:1,000, goat anti-mouse antibodies coupled to Alexa Fluor® 488 at 1:2,000 and streptavidin coupled to Alexa Fluor® 595 (Invitrogen) at 1:4,000. Nuclei were stained with 1 μg/mL Hoechst-33342 (Invitrogen). Airyscan confocal imaging in superresolution mode was carried out using a Zeiss Airyscan LSM 880 microscope equipped with 405-nm, 488-nm and 594-nm laser lines and a 63x Plan APO NA 1.4 oil immersion objective.

For generation of structured illumination microscopy (SIM) images, parasites were stained with 2 μg/ml DAPI in medium for 20 minutes at 37°C. They were then placed on a microscope slide, covered with a coverslip and immediately imaged on a Zeiss Elyra microscope equipped with a Plan-Apochromat 100x/1.46 oil objective. Reconstruction of SIM images was performed in ZEN 2.3 Black software using default settings.

Images were processed using Fiji (Schindelin *et al.*, 2012) and Adobe^**®**^ Photoshop^**®**^ CC 2019 was used for display purposes only.

### Western blot analysis

Immunoblots were performed using saponin-lysed, infected erythrocytes. Parasite proteins were separated on a 12% SDS-PAGE gel as described previously (Heiber and Spielmann, 2014; Bachmann *et al.*, 2015) and transferred to a nitrocellulose membrane (Amersham Protran; 0.45-μm pore nitrocellulose membrane; GE Healthcare) using a Trans-Blot device (Bio-Rad) according to the manufacturer’s instructions. The membranes were blocked with 3% skim milk in TBS for 30 minutes and then probed with mouse anti-GFP (1:1,000, Roche) or rabbit anti-aldolase (Mesén-Ramírez *et al.*, 2016) (1:2,000). The chemiluminescent signal of the horseradish peroxidase-coupled secondary antibodies (Dianova) was visualized using a Chemi Doc XRS imaging system (Bio-Rad) and processed with Image Lab 5.2 software (Bio-Rad).

To perform loading controls and ensure equal loading of parasite material, rabbit anti-aldolase (Mesén-Ramírez *et al.*, 2016) antibodies were used. The corresponding immunoblots were incubated two times in stripping buffer (0.2 M glycine, 50 mM dithiothreitol, 0.05% Tween 20) at 55°C for 1 h and washed 3 times with Tris-buffered saline for 10 min.

### Proximity-dependent Biotin Identification (BioID) and mass spectrometry analysis

The protocol for proximity-dependent biotin identification (BioID)(Roux *et al.*, 2012) in *P. falciparum* was adapted from previous published assays (Khosh-Naucke *et al.*, 2018; Geiger *et al.*, 2020) and mass spectroscopy analysis was performed as described (Geiger *et al.*, 2020). Briefly, 100 mL of highly synchronous *Pf*PhIL1-BirA-GFP or 3D7 parasites at a parasitemia of 7 to 10% were grown until 40 hpi. Then the cultures were grown in culture media supplemented with 150 μM biotin (Sigma-Aldrich) and compound 2 was added for 8 hours. Erythrocytes were lysed with 0.03% saponin and the isolated parasites were washed twice with PBS, taken up in lysis buffer (50 mM Tris-HCl pH 7.5, 500 mM NaCl, 1 mM DTT, 1 mM PMSF, 2x protease inhibitor mix, 1% Triton) and frozen at −80°C. Three freeze-thaw cycles were performed; the samples were centrifuged at 25,000 g for 60 minutes at 4°C and the supernatant was stored at −80°C. For purification of biotinylated proteins, 50 μL streptavidin sepharose (GE Healthcare) was added to the lysate and incubated overnight with end-over-end rotation at 4°C. The beads were washed twice in lysis buffer, once in dH2O, twice in Tris-HCl (pH 8.5) and three times in 100 mM triethylammonium bicarbonate buffer (TEAB) pH 7.5 (Sigma-Aldrich). The washed beads were resuspended in 200 μL 50 mM ammonium bicarbonate (pH 8.3) and on-bead trypsin digest (rolling with 1 μg of trypsin (Roche) for 16 h at 37°C followed by a second trypsin digest with 0.5 μg trypsin for 2 hours) was performed. Next the samples were centrifuged at 2,000 g for 5 minutes, resuspended in 2 x 150 μL 50 mM ammonium bicarbonate (pH 8.3, “first elution: Ambic”), transferred to and collected in a spin column (Pierce Spin Columns with Snap Cap, Thermo Scientific) placed in a low binding tube (Low Protein Binding Microcentrifuge tubes, Thermo Scientific). Subsequently, the left-over biotinylated peptides were eluted from the beads by 2 x 150 μL 80% acetonitrile and 20% trifluoroacetic acid (“second elution: ACN”). Then, the samples were dried using SpeedVac (Thermo Fisher Scientific) and stored at −20°C.

Dried peptides were sent to the Proteomics Core Facility at EMBL Heidelberg. Peptides were dissolved in 1% formic acid/4% acetonitrile, sonicated in the ultrasonic bath for 5 minutes and desalted using an OASIS® HLB μElution Plate (Waters). Cleaned peptides were dissolved in 50 mM HEPES pH 8.5 and labelled with TMT6plex Isobaric Label Reagent (ThermoFisher) according to the manufacturer’s instructions. After labelling, samples were pooled and purified from unreacted TMT label using OASIS® HLB μElution Plate (Waters). An UltiMate 3000 RSLC nano LC system (Dionex) fitted with a trapping cartridge (μ-Precolumn C18 PepMap 100, 5 μm, 300 μm i.d. x 5 mm, 100 Å) and an analytical column (nanoEase™ M/Z HSS T3 column 75 μm x 250 mm C18, 1.8 μm, 100 Å, Waters). Trapping was carried out with a constant flow of solvent A (0.1% formic acid in water) at 30 μL/min onto the trapping column for 6 minutes. Subsequently, peptides were eluted via the analytical column with a constant flow of 0.3 μL/min with increasing percentage of solvent B (0.1% formic acid in acetonitrile) from 2% to 4% in 4 min, from 4% to 8% in 2 min, then 8% to 28% for a further 37 min, and finally from 28% to 40% in another 9 min. The outlet of the analytical column was coupled directly to a QExactive plus (Thermo) mass spectrometer using the proxeon nanoflow source in positive ion mode. The peptides were introduced into the QExactive plus via a Pico-Tip Emitter 360 μm OD x 20 μm ID; 10 μm tip (New Objective) and an applied spray voltage of 2.1 kV. The capillary temperature was set at 275°C. Full mass scan was acquired with mass range 375-1200 m/z in profile mode with resolution of 70,000. The filling time was set at maximum of 10 ms with a limitation of 3×106 ions. Data-dependent acquisition (DDA) was performed with the resolution of the Orbitrap set to 17500, with a fill time of 50 ms and a limitation of 2×10^5^ ions. A normalized collision energy of 32 was applied. Dynamic exclusion time of 30 s was used. The peptide match algorithm was set to ‘preferred’ and charge exclusion ‘unassigned’, charge states 1, 5 - 8 were excluded. MS2 data was acquired in profile mode. IsobarQuant(Franken *et al.*, 2015) and Mascot (v2.2.07) were used to process the acquired data, which was searched against a Uniprot *Plasmodium falciparum* (UP000001450) proteome database containing common contaminants and reversed sequences. The following modifications were included into the search parameters: Carbamidomethyl (C) and TMT10 (K) (fixed modification), Acetyl (Protein N-term), Oxidation (M) and TMT10 (N-term) (variable modifications). For the full scan (MS1) a mass error tolerance of 10 ppm and for MS/MS (MS2) spectra of 0.02 Da was set. Further parameters were set: trypsin as protease with an allowance of maximum two missed cleavages: a minimum peptide length of seven amino acids; at least two unique peptides were required for a protein identification. The false discovery rate on peptide and protein level was set to 0.01.

The mass spectrometry proteomics raw data have been deposited to the ProteomeXchange Consortium via the PRIDE (Vizcaíno *et al.*, 2016) repository with the dataset identifier XXXXXX.

### Growth assay

For growth assays of PhIL1-TGD a flow cytometry assay adapted from previously published assays (Malleret *et al.*, 2011; Jonscher *et al.*, 2019) was performed to measure parasite growth over five days. In order to obtain highly synchronous parasite cultures late schizonts were isolated by percoll gradient(Rivadeneira *et al.*, 1983) and cultured with fresh erythrocytes on a shaker for 4 hours. Afterwards sorbitol synchronization (Lambros and Vanderberg, 1979) was applied in order to remove remaining schizonts resulting in a highly synchronous ring-stage parasite culture with a four hour age window. Next the parasitemia was determined by flow cytometry and the culture diluted to 0.1% parasitemia. Each day parasite cultures were resuspended and 20 μL samples were transferred to an Eppendorf tube. 80 μL RPMI containing Hoechst-33342 and dihydroethidium (DHE) was added to obtain final concentrations of 5 μg/mL and 4.5 μg/mL, respectively. Samples were incubated for 20 min (protected from UV light) at room temperature, and parasitemia was determined using an LSRII flow cytometer by counting 100,000 events using the FACSDiva software (BD Biosciences).

### GlmS-based knockdown

GlmS-based knockdown assay was adapted previously published assays (Prommana *et al.*, 2013; Burda *et al.*, 2020). To induce knockdown, highly synchronous early rings stage parasites were split in two dishes and 2.5 mM glucosamine was added to one of them and parasite growth was measured by flow cytometry over five days as described above. As an additional control, the same amount of glucosamine was also added to 3D7 wildtype parasites. For all analyses, medium was changed daily, and fresh glucosamine were added every day.

Knockdown was quantified by fluorescence live cell microscopy using schizonts 36 h post glucosamine treatment. Parasites with similar size were imaged, and fluorescence was captured with the same acquisition settings to obtain comparable measurements of the fluorescence intensity. Fluorescence intensity (integrated density) was measured with Fiji (Schindelin *et al.*, 2012), and background was subtracted in each image. The data was visualized with Graph Pad Prism version 8.

### Conditional inactivation via knock-sideways

The knock-sideways approach was performed as described previously (Birnbaum *et al.*, 2017). The PF3D7_0308300-2xFKBP-GFP-2xFKBP schizonts were isolated using a percoll gradient(Rivadeneira *et al.*, 1983) and incubated for four hours with fresh erythrocytes. Next, parasites were treated with 5% sorbitol (Lambros and Vanderberg, 1979) to obtain a culture containing rings of a stage window of zero to four hours after invasion. The culture was split into two 2-mL cultures of which one was supplemented with 250 nM rapalog (Clontech). Parasite growth was determined via flow cytometry over five days as described above. Mislocalization of PF3D7_0308300-2xFKBP-GFP-2xFKBP into the nucleus was confirmed by live-cell microscopy 36 hours after rapalog addition.

### Protein schematics

Schematic protein representations were designed using IBS (Liu *et al.*, 2015), predicted protein domains were obtained from plasmodDB (Aurrecoechea *et al.*, 2009) interfered from Interpro (Mitchell *et al.*, 2019) and TMHMM (Sonnhammer *et al.*, 1998).

---

**Figure S1: Localization of PhIL1-BirA-GFP to the IMC in asexual blood stages**

**(A)** Schematic representation of the SLI-based single-crossover homologous recombination approach resulting in a *phil1-birA*-gfp* fusion of the endogenous locus. Dark green, human dihydrofolate dehydrogenase (hDHFR); grey, homology region (HR); light green, green fluorescence protein (GFP) tag; blue, T2A skip peptide; turquoise, BirA* ligase; orange, neomycin resistance cassette; stars indicate stop codons and arrows depict primers used for the integration check PCR. **(B)** PCR analysis of the rendered genomic locus using primers as shown in A using gDNA of 3D7 wild-type and PhIL1-BirA-GFP parasites. Ladder size indicated in base pairs (bp) **(C)** Western blot analysis using mouse anti-GFP antibody detects an approx. 100 kDa protein in *Pf*Phil1-BirA-GFP and no protein in the parental 3D7 cell line. Anti-aldolase antibody was used as loading control. KI = knock-in **(D)** Localization of *Pf*PhIL1-BirA-GFP during asexual blood stage development of *P. falciparum* (stages are indicated) using live microscopy. Scale bars 2 μm. **(E)** Live microscopy of PhIL1-BirA-GFP parasites episomally expressing ALV5-mCherry under control of the late-stage promoter *ama1*. Nuclei were stained with Hoechst-33342. Scale bar, 2 μm. **(F)** Scatterplot showing proteins with a log2FC ≥ 0 in the first elution of both BioID replicates. Green dots are indicating known IMC proteins. Scatterplots showing log2FC values of all detected proteins in first and second elution fractions of both BioID replicates or mean log2FC values of first vs second elution.

**Figure S2: Localization of PhIL1 interaction candidates (PIC) using endogenous GFP tagging**

**(A)** PIC1 (PF3D7_1229300), **(B)** PIC2 (PF3D7_0822900), **(C)** PIC3 (PF3D7_0415800), **(D)** TKL4 **(E)** PIC4 (PF3D7_0308300), **(F)** PIC6 (PF3D7_0530300). Localization of the GFP fusion protein throughout the intraerythrocytic cycle. Nuclei were stained with Hoechst-33342 or DAPI. Scale bar, 2 μm **(G)** PCR-based confirmation of the correct insertion of TKL4-GFP-encoding integration plasmid into the targeted loci. Ladder size indicated in base pairs (bp). gDNA from parental 3D7 was used as control. KI = knock-in.

**Figure S3: Localization of apical annuli proteins using endogenous GFP tagging**

Localization of the GFP fusion protein **(A)** *Pf*AAP6 (PF3D7_0508900), **(B)** *Pf*AAP4 (PF3D7_1318700), **(C)** *Pf*AAP2 (PF3D7_131280) throughout the intraerythrocytic cycle and in gametocytes **(D)**-**(F)**. Nuclei were stained with Hoechst-33342. Scale bar, 2 μm.

**Figure S4: Alignments of** *Tg***AAMT/***Pf***AAMT and** *Tg***AAP2/** *Pf***AAP2**

Alignment on amino acid level of (A) *Tg*AAMT (TGGT1_310070) with *Pf*AAMT (PF3D7_1455200) and (B) *Tg*AAP2 (TGGT1_295850) with *Pf*AAP2 (Pf3D7_1312800)

**Figure S5: Conditional knock-down / knock-sideways of PICs and AAP proteins**

Live cell microscopy of **(A)** PIC1-GFP or **(B)** PIC3-GFP parasites 36 hours after treatment without (control) or with 2.5mM glucosamine. Growth of parasites treated without (control) or with 2.5mM glucosamine determined by flow cytometry is shown from three (PIC1) or four (PIC3) independent growth experiments. Quantification of knockdown by measuring intensity density and size (area) of parasites 36 hours after treatment, displayed as individual data points of control (blue) and 2.5mM glucosamine (red) parasites with mean +/− SD of two independent experiments. N indicates the number of measured cells. **(C)** Live cell microscopy of PIC4-2xFKBP-GFP-2xFKBP parasites expressing the 1xNLS-FRB-mCherry mislocalizer under the control of the *nmd3* promoter untreated (control) or rapalog treated (+rapalog) at 40h post addition of rapalog. Growth of parasites treated without (control) or with rapalog determined by flow cytometry shown from three independent growth experiments. Live cell microscopy of **(D)** *Pf*AAP6-GFP, **(E)** *Pf*AAP4-GFP or **(F)** PIC2-GFP parasites 36 hours after treatment without (control) or with 2.5mM glucosamine. Nuclei were stained with Hoechst-33342. Scale bar, 2 μm.

**Figure S6: Targeted gene disruption of PIC4 and PhIL1**

**(A)** Schematic representation of the SLI-based single-crossover homologous recombination approach resulting in a truncation and GFP fusion of the target gene. Dark green, human dihydrofolate dehydrogenase (hDHFR); grey, homology region (HR); light green, green fluorescence protein (GFP) tag; blue, T2A skip peptide; orange, neomycin resistance cassette; stars indicate stop codons and arrows depict primers used for the integration check PCR. **(B)** Live cell microscopy of PIC4-TGD parasites during asexual blood stage development (stages are indicated). Nuclei were stained with DAPI. Scale bar, 2 μm. **(C)** PCR analysis of the rendered genomic locus using primers as shown in Figure S2A using gDNA of 3D7 wild-type and PIC4-TGD parasites. Ladder size indicated in base pairs (bp). Schematic representation of the 338 aa full length PIC4 and the truncated 93 aa PIC4-TGD fragment. **(D)** Growth curves of PhIL1-TGD vs. 3D7 parasites monitored over five days by flow cytometry. Six independent growth experiments were performed, and a summary is shown as mean +/− SD percentage of growth compared to 3D7 control parasites after 2 parasite replication cycles. **(E)** Live cell microscopy of PhIL1-TGD parasites. Nuclei were stained with Hoechst-33342. Scale bar, 2 μm. **(F)** PCR analysis of the rendered genomic locus using primers as shown in A and gDNA of 3D7 wild-type and PhIL1-TGD parasites. Ladder size indicated in base pairs (bp). Schematic representation of the 225 aa full length PhIL1 and the truncated 117 aa PhIL1-TGD fragment.

**Table S1: BioID list** Complete list with log_2_ fold change values of all detected proteins, non-detected proteins (NA) were replaced by 0. IMC proteins are highlighted in green, proteins marked in blue were selected for further validation.

**Table S2: Oligonucleotides used in this study**

Restriction sites are highlighted in red.

