## Supplementary Figures for "Identification of novel inner membrane complex and apical annuli proteins of the malaria parasite *Plasmodium falciparum*"

### Figure S1

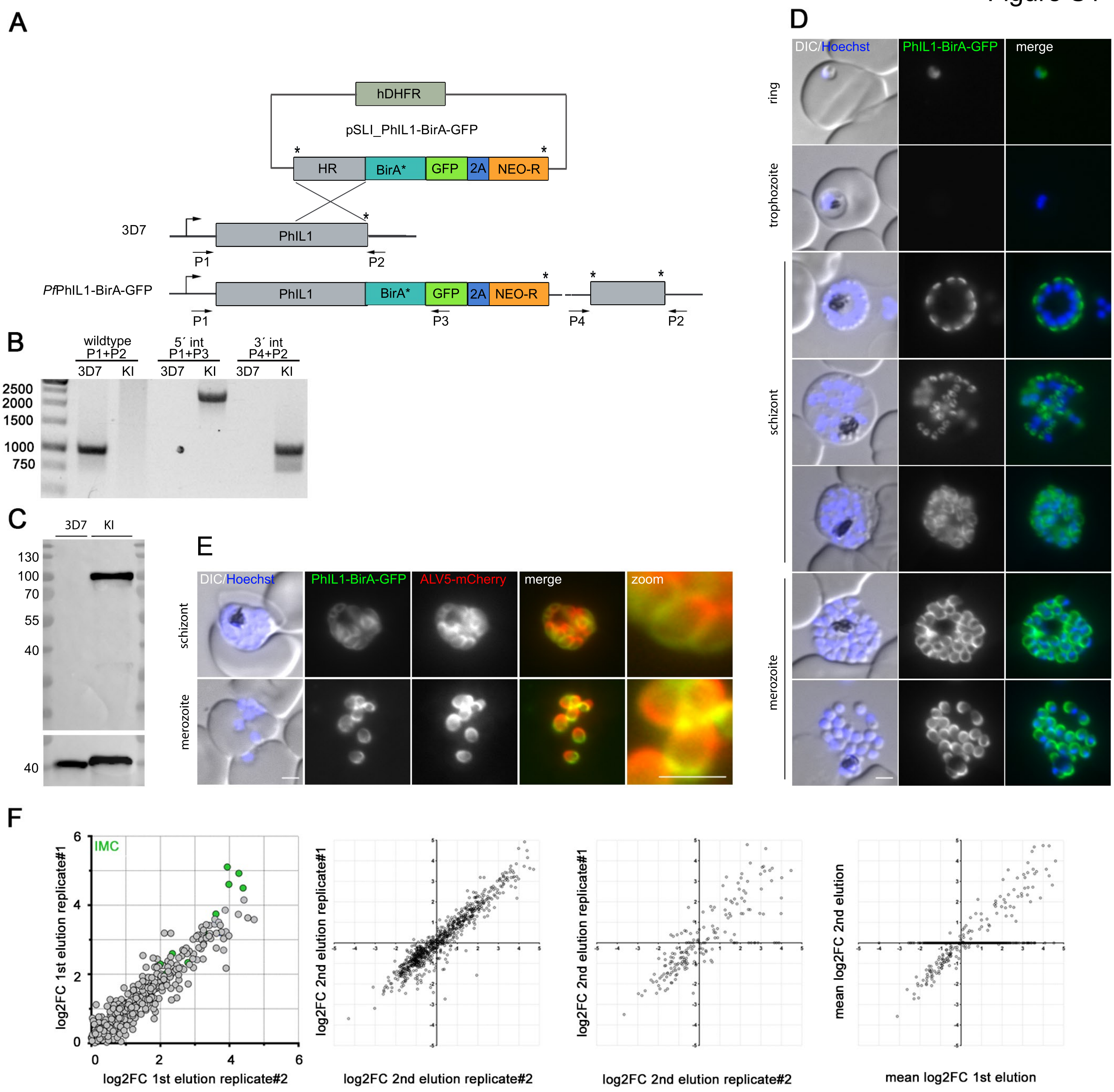
A
hDHFR
pSLI_PhIL1-BirA-GFP
HR
BirA*
GFP
2A
NEO-R
3D7
PhIL1
P1
P2
PfPhIL1-BirA-GFP
P3
P4
B
wildtype P1+P2
5´ int P1+P3
3´ int P4+P2
3D7
KI
2500
2000
1500
1000
750
C
130
100
70
55
40
D
DIC/Hoechst
PhIL1-BirA-GFP
merge
ring
trophozoite
schizont
merozoite
E
ALV5-mCherry
zoom
F
IMC
log2FC 1st elution replicate#1
log2FC 1st elution replicate#2
log2FC 2nd elution replicate#1
log2FC 2nd elution replicate#2
mean log2FC 2nd elution
mean log2FC 1st elution

Figure S2

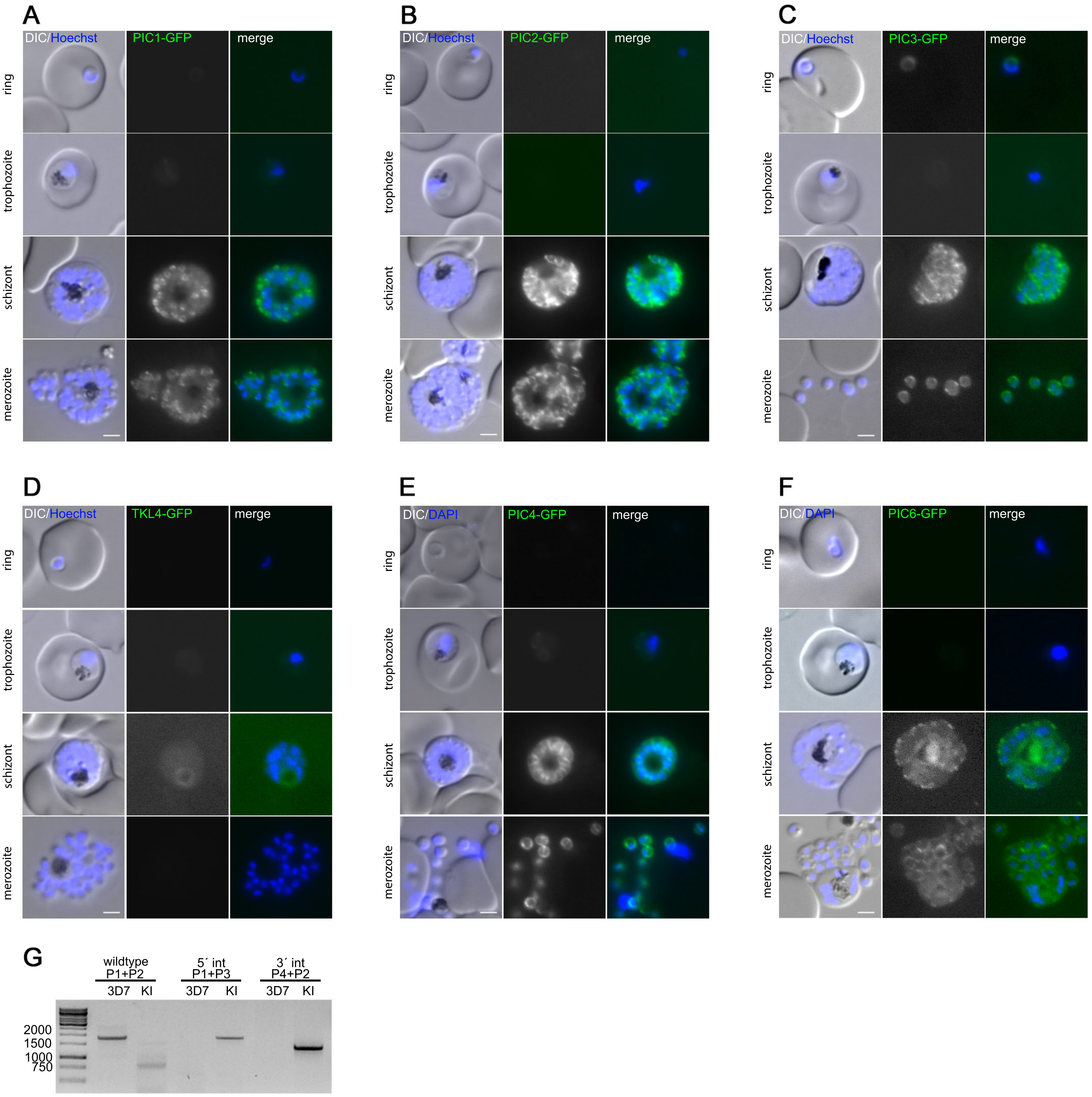
A
DIC/Hoechst
PIC1-GFP
merge
B
DIC/Hoechst
PIC2-GFP
merge
C
DIC/Hoechst
PIC3-GFP
merge
D
DIC/Hoechst
TKL4-GFP
merge
E
DIC/DAPI
PIC4-GFP
merge
F
DIC/DAPI
PIC6-GFP
merge
ring
trophozoite
schizont
merozoite
G
wildtype
P1+P2
5´ int
P1+P3
3´ int
P4+P2
3D7
KI
2000
1500
1000
750

Figure S3

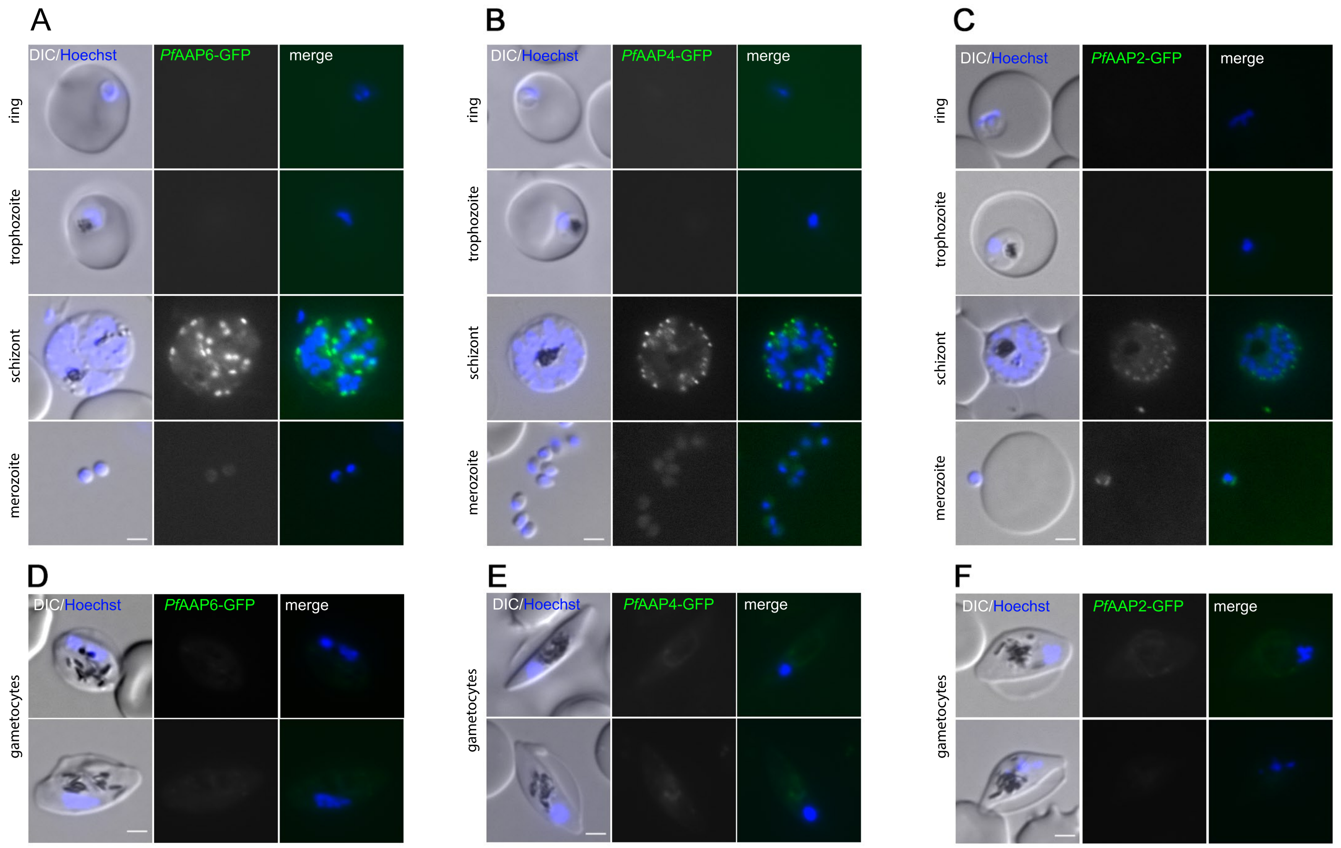
A
B
C
D
E
F
DIC/Hoechst
PfAAP6-GFP
PfAAP4-GFP
PfAAP2-GFP
merge
ring
trophozoite
schizont
merozoite
gametocytes

Figure S4

## A

```
Alignment of Sequence_1:  [TGGT1_310070.xprt] with  Sequence_2: [PF3D7_1455200.xprt]

Similarity : 140/204 (68,63 %)

Seq_1  1     MTA-YGKVDYWDERYKRDVEPFDWFQRYAGLKPILLEAGLQASSRILVLGCGTSRVSEEM  59
             | | |||:: :: : : ::|:::   : :  :  :: :::: ::: : :   :::  |:
Seq_2  1     M-AVYGKIS-YWNERYTNEEEQFDWHQRWYGVKHIFTELEIKNDANILNIGCGTSKFSEE  58

Seq_1  60    YADGYRKIVNVDYSNVCISHMQRRCADKEEMTFLHMNALDMKQLDDGDFDLVFDKGTMDC  119
             :|:   : ::::: ::: ::   ::: :: :::   ::::  : :::::::: ::::: ::
Seq_2  59    MLDSGYTNITNIDASSVCIKKMQELYNDKPNLKYILMNVCDMREFTNEEFDLIIDKACLD  118

Seq_1  120   VLCGDNSFDNVQKMLREVSRILAPGGVYIVVSYGQPNFRLSHLQREEYGWSVTMKTIQKP  179
             ::: ::: ::: :: : : ::: :: :::::|:::  ::   :: :: |: : :: ::  :
Seq_2  119   SIVCSEDSLKNVEEMLSEVSRILKSNGIFVIISHAQPAYRLVYLQKEDYNWDITVKTVQR  178

Seq_1  180   SINVQAPIDEKDNVHYVYICKKAASEAAQAPAQVSSDAAKGAPTN  224
             : :::: :: : :|:||:|||||| :
Seq_2  179   PMLGIVAPPVDDNLHYIYICKKKHTSK------------------  205
```

## B

```
Alignment of Sequence_1:  [TGGT1_295850.xprt] with  Sequence_2: [Pf3D7_1312800.xprt]

Similarity : 1591/2283 (69,69 %)

Seq_1  1     --------------------------------------------MTSPSSPLPAPSAPS  15
                                                         : : |:    :: :
Seq_2  1     MSFLLALNESKKSAEGSVASDITEEVFEKEIHEMNSLNMIMHIEDLRDKSFVDVKEKKDE  60

Seq_1  16    VPYALHYSSTPQDPICSTPSFPSFSTSSVWGNPFT-TEH--------KAPQVKGDHLS-P  65
             : : ::: :: ::::::: : :: :  | :| |  :::         |   :| || :
Seq_2  61    NKIKIKNTDEKEKEENGEKEGEEKGG-S-IKNSFMKIFQKKKKKKKKTKKIK-DHNINN  117

Seq_1  66    ASARPFLGTVRSISPAPQDAERSLASRGHFLPEKRPE-QNGWQLAAGAGHQSRPD-VASL  123
             ::::  |:|: : : : |  |    :| | :::|  :: |   | ::   : ::|:::| :::
Seq_2  118   IYDSNCLLTN-PDNVDP-D-KYYL-S-FNELKKENNENVN-EENYSSKDKQKNIDNSKDI  171

Seq_1  124   THALSGSQTE-ESPRTEAALAGEVR-SAGSRVLEHVVASPVKEASIRVSQEAPDVSL--K  179
              :::::: :| :: : :   ::::  |  | :::|:|:: :: ::|:::  |:::   |
Seq_2  172   VQNEENMNKEQIKDNMKRKKKKIKLSSYGTNSIENVKMKKYDSNKINNMDETYNNIYKKK  231

Seq_1  180   KSRTLAERFFLISSPPKAP-KAETRS-AQDNEEGIKGKKGEGRDACGFGAQRAPAEERGR  237
             :|::|  :   | : :|| |  :|:|: |  :  :: : | :::: | ::: ::   :::|
Seq_2  232   QSNDLK-DKKLKTKRFKNLKKKIT-SPKFTMSTKTKNKKN-G-NINENNFNFDPDDDLKK  287

Seq_1  238   DWASDGETIAEREEGLDLTGGDDHVGTSSQQVKQ-SSFGEEPSDPATSWERGESAASLE-  295
             ::::::::| ||:   :::::::::::::::| :::::: :::|::::::: :::::::|
Seq_2  288   EIEKIKIDI-ERIK-NKGNIKRMYDDLKYILLFQNEIVDIIIEDIRKNKVR-TKYEMLED  344

Seq_1  296   SLFRGRSLQQKKKRWWAGLASSLGRRSRSVECGRRGGDAAEGRGAEESEKGRLPQDTPGS  355
             |   ::::   :::::::|::: :: :::|::: :: ::: :::   :  :|::|
Seq_2  345   STVKDDVNKVILNHITFDLVKFKYINTKLKELILCVYDTKKFNG-LVKRIVNNKAHEMRK  403

Seq_1  356   RSGPQSAAELRKETREGRGVRDEPGENESDLWRRPASPEEDVARKERRREKRER-ETLDE  414
             ::: :::  ::| ::::::::  :| |:::  | |:::: :: ::::|| :
Seq_2  404   IVDKNGELTVAK-SMLSFLYKQLKYCIE-DEIQCYYS-EVKKLNDDIHNIKRNVMNIIFK  460

Seq_1  415   SSSGFLPMTELLKGQ-GE-G-RAGRTSDSSFVSSRASTKKSVTTKLSRSLSRLFGGKKKK  471
               |: :  |  : :  |  |  :::|:::::::::::::::|: ::: ::: :: ::|
Seq_2  461   --SINDVCTMPDPYQIYEMGDKINRKKGTKENYNLNDINDDEQIK-CEQNFVEYER-KQQ  516

Seq_1  472   TKGKERAGEQADGTS-EGRRHSADGGPRQAHGRRDSSTSVSSLSSRSGPSTD-LSRSSSP  529
              : :  : :::::|  :::|::::: : :::::::::: ::::::  ::| :::::
Seq_2  517   MICPDYSSDLNLFFSLYDNIHIERKKYIKNKSVMFKNYCTDVDNDDDHNINDNINHNNNV  576

Seq_1  530   CSPCFSASQWEEAFSSLPPIPPSVAALSDT-CQDLLLDQLKVLERGVQQLLAGQGSDVVL  588
             ::  : :::    ::   |: :: |::| : :: :: :::   ::::   ::  ::::  |
Seq_2  577   NIYGNMKNIKMSTYRVNNNILDKD-AQFVTSSKNINESYIPNYTNKSIRSVVSQ-NY-VS  633

Seq_1  589   ACIADEPRISEVMKPNQPSGSA-YRSQLLVDEKGHDEGGDLSRQGAVEGACTGERREGNS  647
             :: ::|  | : :::::  :::|: |::  ::: :::: ::::|:: ::: :|:: :::|:
Seq_2  634   PFNMIENRHKTDDFIYGDRNSNYYINKSYEKTSNINNNIPLNVKFLNKNAVNANYNVSNM  693

Seq_1  648   RGGGELGEGEASLFFSPERAQKVCRRSQIPRAVGRLREELTAEHTASIQLLTDERNRLVL  707
              : ::|  ::  ::   : :::::: : :   |   ::: : :|:: ::::|   :::::|:
Seq_2  694   KMKKEHYLDDLKNYYCNINIPWIGKTCTNP-SSSMSKNILKRKSNNKI-KGRVIDAPLFH  751

Seq_1  708   NKLLVGIHEVRGLVRTFTDVSRDAQGEAAEI-LADAAQLRAWSAK-HGELAEGRDLVEHY  765
             : : ::| : :    :::::::: : :: | : :   :::::| : : :   : ::|:
Seq_2  752   SAPDNDI-NHTNDTYDINNIGFGNDIDNNNIFEYVGYKNSDIYNKNIYPNNDAYSNYQHD  810

Seq_1  766   HRQVAESANGVIREYH-RRCIQLNSDIQLMRDQISQVVHAAFLEASRLPPLHSLYASSSE  824
             :: | :::|:::: :| : :    : :|   :::  : :::::  ::::   | :: :::::
Seq_2  811   SFNV-PYPNELFNKDHVMYNNNHN-MIHDENLVYNYDYKINDAQRNNY-P-PNNVHNNHM  866

Seq_1  825   RNCVDAFLSSCPYSGDPRIFMVSPPFLSRLQLYALSHLPPSRSAHAPDSPAES-PTGAVA  883
             ||:: : :::   :::| | | ::    ::: :: : ::   ::: :: :: :    :::
Seq_2  867   RNQM-NNIYIDKDMLDGR-F-HKMSHIINNIDEKCYDGYNYKYYNLGENHNNSLNMYHNM  923

Seq_1  884   SSTASSLHVGRHAVVVPVSSSALFQPGR-CVQSVPPGLLLRSSKEDLAALEHRLEGKWGS  942
             ::::: ::: :: :: :   |:::  :  :|  : ::   |::::|||||:  ||::| :||||
Seq_2  924   NGNIYTERFAKQSNGTTVRCFKSEVNIRNKDTNKYSGFEIISSKEDIKKLENLLNDKWGS  983

Seq_1  943   FNLFEAWQNDAMKWQWRDAMMQRNFATREEEIERWRRDSEARVLAAVDLRLQELWAGRMQ  1002
             |:|||  |::| :||||:|: :|:: ::: :|:|::::::   ::  :: :: ||| |||
Seq_2  984   FKLFEKWIKDSSKWQWIDLKLQKEYINSNIELEKLKEEKTKYLEKCINNKIKELWCGRMK  1043

Seq_1  1003  ELRRVAGENLEDSFVSLLVDRVPSVADVVQRVQQKRRATGSASLSSASSLSCFHFASALA  1062
              |::: :  | |::::: ::: :::::::: ::: :: ::|:    :::|:  :: :: :: : :
Seq_2  1044  GLMNLKLCNMERKILKRFSKYHINQDIYDVVLDFERNSMCYQDLYAKLLNIEVYSKKVDF  1103

Seq_1  1063  AGVSNEEVEAAFLAAEDMRSRLVEVASSEVFRQLLHAQQIVADYREGKSREKLLGVLEET  1122
             ::::|| :|::      :: :  : : :: :: :::  :::   : :::::|: |  :: : :
Seq_2  1104  INKYNEYNEQDAGGDYIYKYNKDKKKVNDENNKISKKGYVQSLIYSFNKKSQKSIN  1163

Seq_1  1123  VGKQMREICHLRTRLEEESDLACQPDRIRDECQKTEAAITMLEEELIAMQEKRTLRQQRL  1182
             ::: ::: ::   :::   ::  :| :: :: :: ::: :   :  ::: :::: :: :
Seq_2  1164  KNNVQIQKEKKYNSIMKDPTTSVQYKNDHIYNIMDISNNRANAVINKIETINNKADINNN  1223

Seq_1  1183  LDLRDSAQTKLQRVTALFDDDQALCRSLEEQCAGERRRGERVERLWQEIGIEKLEEMKQE  1242
              :::: ::|:  :::: ::|: :: ::    :  :::: :: ::: ::: :: :: :: ::
Seq_2  1224  KKEIHTNLTENTNKKNNLGDNNRIDEDTTKKDQTYDKGKKEQKNEPKQEQKKEPKQEPKQ  1283

Seq_1  1243  LARLKGENVALQADLEVARFELDVKRKKEELKREEAQRRQAEAARKQQEMREKKGERERK  1302
             :::::: ::: |:   :::  |::::|:|: ::: : :::   : |::: ::: :::|
Seq_2  1284  EPKQEQKYEQKQEQKHEQKHEQKHEQKHEQKIDQKLMKKPKNIYSNAQLNNMTHIGSFSI  1343

Seq_1  1303  KRALKERDVAERDDAAETQSQRKSPLDSSGLIVRWHKATSSPLPTGPGSQAGNEGEKKHE  1362
             |:: : ::|: ::|: :    | |:|: ||:|::::: :| ::: :: : ::: : :
Seq_2  1344  KKNKSHFDISLQDSGGLVNSKRNSFNDSFSLFRSECNKRISDISLSSLKIKTRDINNEED  1403

Seq_1  1363  REAYGVEEWEADRASDSQGMPFSRASTSKRVGHFSSLFKWPSSKRSSSTPAELRTREGES  1422
              : :::: ::|||::::::::  :::::::: :: :  :::::::::::  ::: :: ::
Seq_2  1404  ISFESSKIDEAEGEEYGEDGDNNDDENIEKTKRTKSNLSTFRRLFNLKRKKKDKEEKPKK  1463

Seq_1  1423  GCEDAESPNGEGPHLRATAIQAASQRPVDSGVGEAHKTSRAAPEVDGLKSLSSSAGKNAE  1482
             :  ::: :: : :: :: ::  :: :::: :::::::: : |   ::: ::::: :::: :
Seq_2  1464  SDVFPSESFSLSYFKRQPSSDNANNSMIHKKENVHNLDIDNNMDDDNNKSSVVLLNNKSLS  1523

Seq_1  1483  GPAEGGEASGRGGLVCADSFSRGIVAGEQTRESRENSGYEAKTGTAELLSPSPNSVLIRG  1542
             :: :  ::::::::::: ::::::: :: :: ::::|:::::    : :::  :  :: :::   ::
Seq_2  1524  VEGYIYDNESIIEKEVGEHLHIENMEDKHISLDDINNEEIYDSHSKSSINHNIYLNDIN  1583

Seq_1  1543  SSAPPPGQDPRESRRERDSGLSGTRGTDHVRDVQPSSPPPEKAPGKSALRRFFSGSPEWA  1602
             :::: ::|::: :: ::::::| :::::  ::: :::  :  : : : :: :   :|:  :|
Seq_2  1584  KINQIMEQTEHKNYAHNIDLIKGVKNKPFKADDKNTIKEKNNKITEKVTDLNDSYSFLNS  1643

Seq_1  1603  AGLRRRPSSKKQDEGDEVSADRSGQKAGCAAAAQQAVTEDLSGGGGRGTALTTAHTQGDP  1662
               : ::::|::  :: :: :::: :::   ::: :::  :  |::::::: :: :: ::|:
Seq_2  1644  SRSSDEENDKGECKQSNEMSNIDNKHDNTGNDRNEHNGDYKSNPESNNKNNLKSNNKSDQ  1703

Seq_1  1663  GRGDQPTRASSLFFRPESWSETKTRRDECARGRVSPPRHRSGEQSQSCFSTPDALDVVRH  1722
             :::  :: :: :: :: :|:::::: ::::::: ::::::: ::|:  ||:::|::::::::
Seq_2  1704  NNNIRNYSKNNSKSDNKNDSINDKIESKENNNKPCYKKQSSSHRSQKSQSIIEDELKKHK  1763

Seq_1  1723  AEVQAKFARGQEARRKEERARELLGMRSSAASRGDSTKELSRDSSGNQDVEHRRQRNELP  1782
             :|: :: :|: : ::| :::: :|:::: :::::  ::::: :  |::|: : :: :: |:
Seq_2  1764  FEDHVINIRRKLSKEKMIDDNIDEGSIKNTSRKKENSLSRSDVSYHSKEEGNKYEKYSNL  1823

Seq_1  1783  REGCVGWGQRTVGKEEPFSQDDETISGGMSHPEGAERGAYPSDGPHSGDTSLSLHADRDS  1842
             ::: :: ::: :  :::::  ::  : :: ::  : : : ::::||::: :: :::::
Seq_2  1824  SFFKKTDKDNKINKFFKNLGRFSFTTSKKKNNKSVQGIYTNDIILHSFSDESDLEEGYEQ  1883

Seq_1  1843  EVQRRTPSKEEQSKANVLSPSSPLWQKHAALCSALGRLERTETSSARGRSPKAGGLDQKS  1902
              :  : ::|::: :: : |:  : ::| :: :: : :::::: |::::: |
Seq_2  1884  KNKTYNNDSEKIYSKSFVLNSYVGKKDEASNINTSEQVDNLYSYINKLKKKKDSKELDKT  1943

Seq_1  1903  ESDMDVLEEETTQKETENVGSGSSEARPTSECRDSGEAQAREGNSSVESEASHTFLPFVS  1962
             ::|::|:     :|  :::::: :: ::: ::::  ::|::::: : :::   :::
Seq_2  1944  QNDESVIKDNGSKVEVKKRKEKDDKDDDKDDDDEDDDKDDDEDDDEDDDKDDDGDDDEDD  2003

Seq_1  1963  SSSRTSPEEGLQLFFCQCRSSDRDQQSSDSRKRRAWEVEDTATPRALQEAREGSRVLEER  2022
             :::::: |:   :   :: :: :::  ::|:::::::: : :: : :: ::   ::: ::|:
Seq_2  2004  DEDDDDGEDDDYNNNGDNTRINSKKEDNKISQNNYKHSNSNNLHDKNEKKDRKNNPED  2063

Seq_1  2023  AKDGNGNVPPLDLRGGNAGGTTERRSTFVSLEGIGLSRGSEHGKQNRGESVERLPTTGFP  2082
             ::: |:|:   : |   ::: ::|::: :::  : : : : : :: : :: : : : :
Seq_2  2064  DHITNNNYSNDVNRYAENKYNNENILSPRFGYSTSNNDAFKNADKLSESNSSKDEISEYS  2123

Seq_1  2083  GSDGPAPTCGIKGDARAGHTDNFSLPSGSSLHLVGPDGLCLFGRSSSSVVSAWENRGAKR  2142
             :::: : : : ::::: :::::   : ::|: : :: |:     ::::::::: : ::::::
Seq_2  2124  LMENKNSKKDNGEENKGKVQFFKAQKDDSNNIMDKKDKKKENIKEKNEETGKNKSSRSS  2183

Seq_1  2143  DSGDRGNSFSGAEAASRPSWEAKEQGKGLTSESLVVGERQIPGLPTVAQFSNSGNAHDAP  2202
             ::::::::::: :::: :: :  : :: ::|:::: :   :  :: : || :::::
Seq_2  2184  SNSSLLDNINRDMNINHKHESSDNEYDDDDKEIIKLSDDKTDNKKISSFHSNNYDHEYNK  2243

Seq_1  2203  SDSNVSGSARRSASAQTVAKRSPRPTTSTDSERQFDCLGSIIEAGLDSGGGHSESSVPED  2262
             :: ::::|  :::::     ::| : : :::: :  ::::::  :: ::::::: ::: |
Seq_2  2244  KNAKSGISSNFIEDKKSTSIFSKKSKMREENSSNSVEIIRLSSFKSRNDLKKKKIGGNND  2303

Seq_1  2263  IGNVTTHREKATGETVPQASSLQTTGFPSWSEIPGGPPVASCSSLRRTAGDNVPGGLPGE  2322
              :::: :: : :| :::  ::    :   ::|::|:|:::: |: :   :::::::
Seq_2  2304  KDDSLSCDNISNGYLSETKKDDYSYYNYVQSILPFGILGNKGSNDSYSSNSIKEDEKT--  2361

Seq_1  2323  EQLSDEKRSSFVSSAPAGVSLSLLGSFGEGDGAQGQGNTAGEGRREHEPCGGLPRPRGGP  2382
Seq_2  2362  ------------------------------------------------------------  2361

Seq_1  2383  EALSCDAPKAIFFSLSATESRVGEQNGETKQKEQVGPSSSVSAHHTVDAGEEEKAADPED  2442
Seq_2  2362  ------------------------------------------------------------  2361

Seq_1  2443  LAGGNQHLIAGSVRIAGGCLRPRPFPADKLRFGELDEKGEDKDRKPETSPPRAGGGNEGK  2502
Seq_2  2362  ------------------------------------------------------------  2361

Seq_1  2503  GSFPPFRSSLSRSFSCDSSLFSVPLTDERSSASRSPPASSPGFERDGYTAVHTPETEEDQ  2562
Seq_2  2362  ------------------------------------------------------------  2361

Seq_1  2563  QASPPPNQTTLRESVTVEPGEDDPMSFLLPSPRAALEPVGCSPSSSEREAEVRERGSDGA  2622
Seq_2  2362  ------------------------------------------------------------  2361

Seq_1  2623  GKDSKRRRRTEFPFSASAPFSFRRSLDEGKALFRGSTFFGRKGGGREEKEATAGLLPRAG  2682
Seq_2  2362  ------------------------------------------------------------  2361

Seq_1  2683  SLDFSILSRRGNKKVCSFGILGSVALDTWRDEEGGLVTEYSPIETFRFNLTQTSHETGQV  2742
Seq_2  2362  ------------------------------------------------------------  2361

Seq_1  2743  DVRNAVLNSPVFALFDVDEKRQILELLISSITHVTEECTIFHRGEKVDVLYFVERGELEM  2802
Seq_2  2362  ------------------------------------------------------------  2361

Seq_1  2803  AAVGPDPENHAEEANLGEREESAVGSAPDSQEAELGNHRHSAMAFSVLQIPSGAFVLPRA  2862
Seq_2  2362  ------------------------------------------------------------  2361

Seq_1  2863  FLRAGCTAYRVRAVKPSSLYKLTAQSFQTVANGAVLRRASDLLAYMLRCPILEALTKREA  2922
Seq_2  2362  ------------------------------------------------------------  2361

Seq_1  2923  SQLIPLMRWQQFLPNEIILHQEQISRTMLIVMKGKARGVRRLSRHGRPETLEYYEEGSCI  2982
Seq_2  2362  ------------------------------------------------------------  2361

Seq_1  2983  NYLCLLQELPNSTSVIAEQPEGCIVASLSASDFVGLMGSAERLLDRGQETADGGKGFSGR  3042
Seq_2  2362  ------------------------------------------------------------  2361

Seq_1  3043  IDLGEVKKTWHKVLKMAGPR  3062
Seq_2  2362  --------------------  2361
```

Figure S5

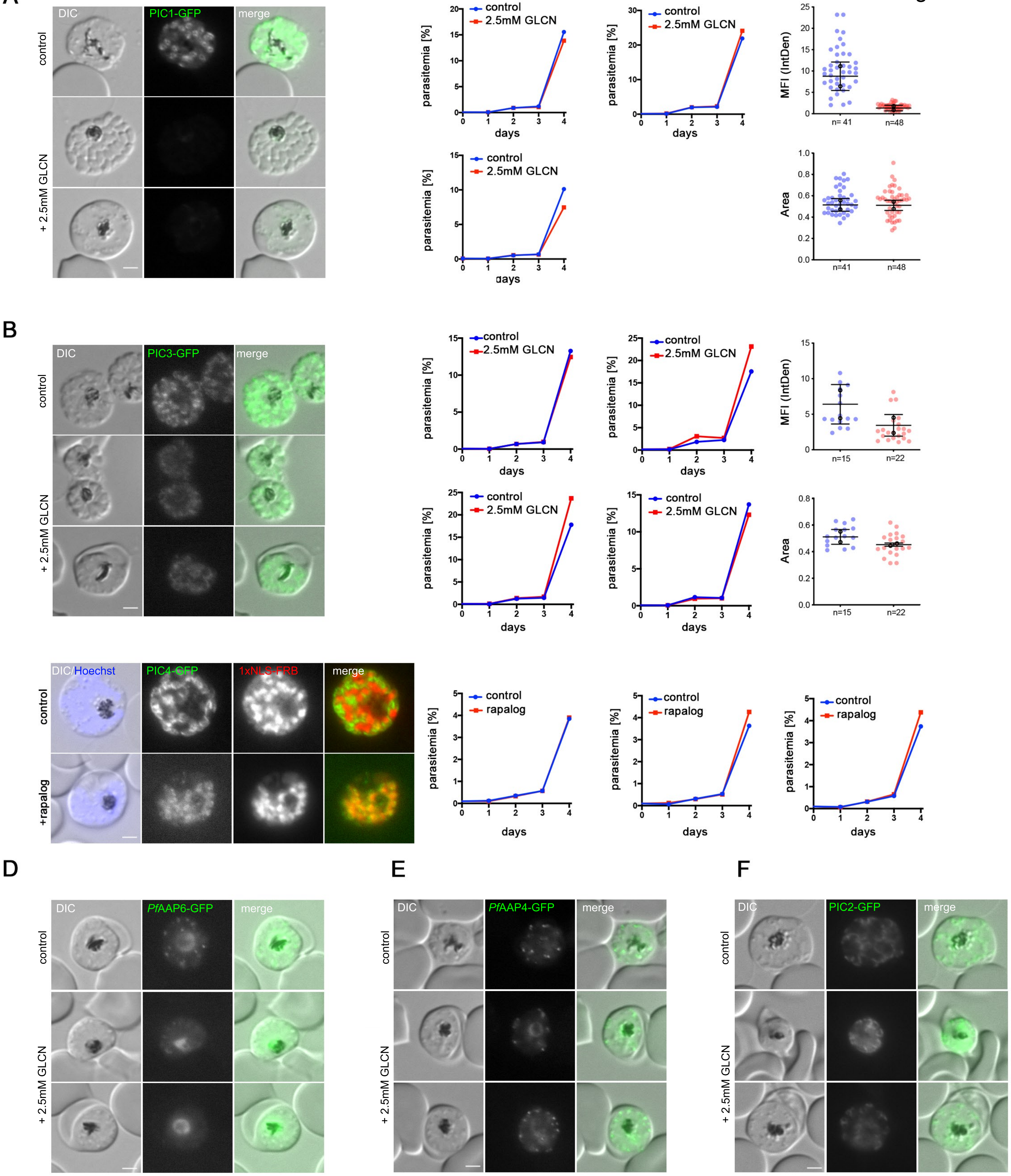
A
DIC
PIC1-GFP
merge
control
+ 2.5mM GLCN
control
2.5mM GLCN
parasitemia [%]
days
MFI (IntDen)
n= 41
n=48
Area
n=41
n=48
B
DIC
PIC3-GFP
merge
control
+ 2.5mM GLCN
MFI (IntDen)
n=15
n=22
Area
DIC/Hoechst
PIC4-GFP
1xNLS-FRB
merge
+rapalog
rapalog
D
PfAAP6-GFP
E
PfAAP4-GFP
F
PIC2-GFP

Figure S6

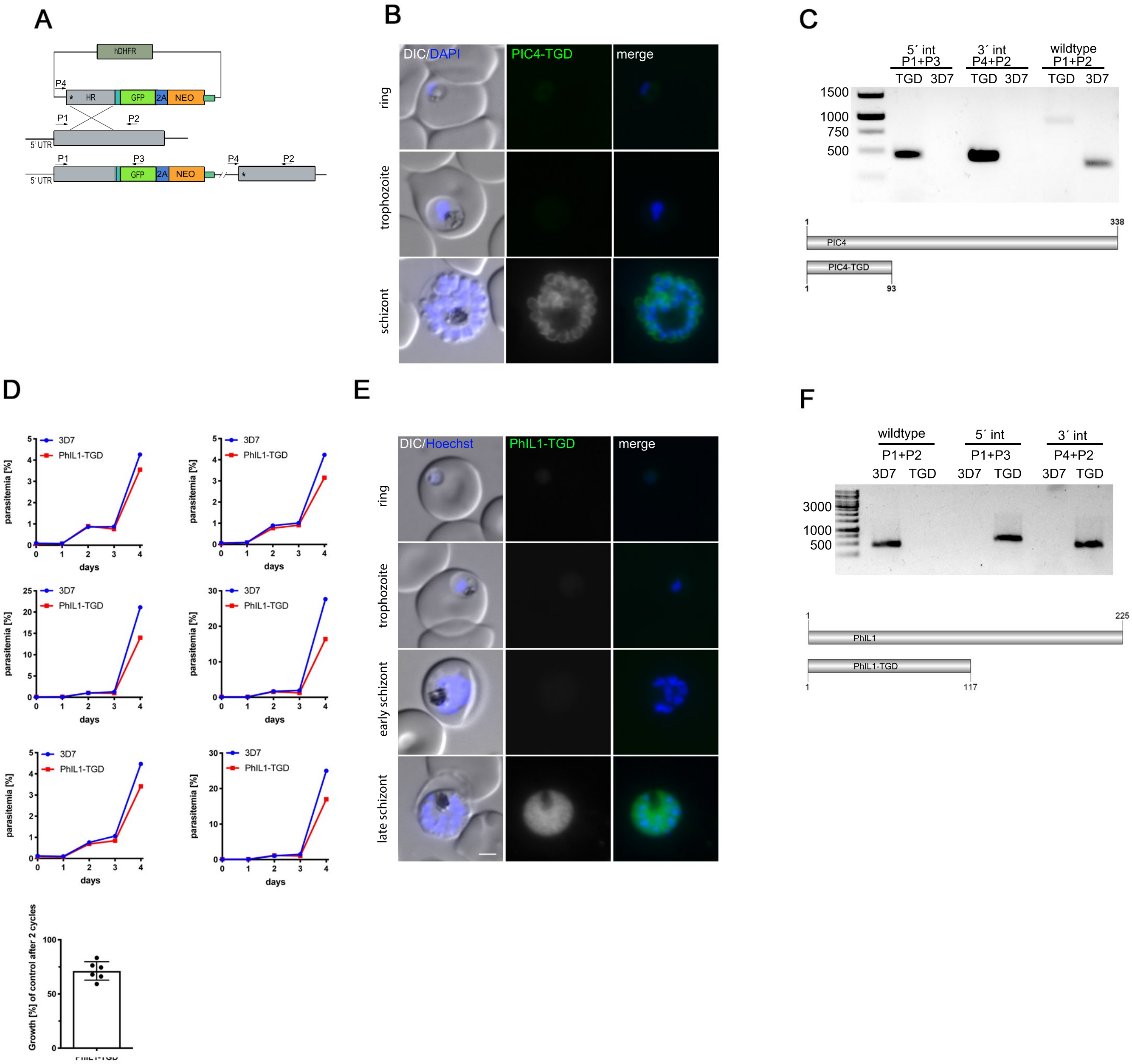
A
hDHFR
P4
HR
GFP
2A
NEO
P1
P2
5´UTR
P3
B
DIC/DAPI
PIC4-TGD
merge
ring
trophozoite
schizont
C
5´ int
P1+P3
3´ int
P4+P2
wildtype
P1+P2
TGD
3D7
1500
1000
750
500
PIC4
PIC4-TGD
1
338
93
D
3D7
PhIL1-TGD
parasitemia [%]
days
Growth [%] of control after 2 cycles
PhIL1-TGD
E
DIC/Hoechst
PhIL1-TGD
merge
ring
trophozoite
early schizont
late schizont
F
wildtype
5´ int
3´ int
P1+P2
P1+P3
P4+P2
3D7
TGD
3000
1000
500
PhIL1
PhIL1-TGD
1
225
117
